# Chromatin-immunoprecipitation reveals the PnPf2 transcriptional network controlling effector-mediated virulence in a fungal pathogen of wheat

**DOI:** 10.1101/2022.06.16.496517

**Authors:** Evan John, Karam B. Singh, Richard P. Oliver, Jessica L. Soyer, Jordi Muria-Gonzalez, Daniel Soo, Silke Jacques, Kar-Chun Tan

## Abstract

The regulation of virulence in plant-pathogenic fungi has emerged as a key area of importance underlying host infections. Recent work has highlighted the role of transcription factors (TFs) that mediate the expression of virulence-associated genes. A prominent example is Pf2, a member of the Zn_2_Cys_6_ family of fungal TFs, where orthologues regulate the expression of genes linked to parasitism in several plant-pathogen lineages. These include PnPf2 which controls effector-gene expression in *Parastagonospora nodorum*, thereby determining the outcome of effector-triggered susceptibility on its host, wheat. PnPf2 is a promising target for disease suppression but the genomic targets, or whether other are regulators involved, remain unknown. This study used chromatin immunoprecipitation (ChIP-seq) and a mutagenesis analysis to investigate these components. Two distinct binding motifs connected to positive gene-regulation were characterised and genes directly targeted by PnPf2 were identified. These included genes encoding major effectors and other components associated with the *P. nodorum* pathogenic lifestyle, such as carbohydrate-active enzymes and nutrient assimilators. This supports a direct involvement of PnPf2 in coordinating virulence on wheat. Other TFs were also prominent PnPf2 targets, suggesting it also operates within a transcriptional network. Several TFs were therefore functionally investigated in connection to fungal virulence. Distinct metabolic and developmental roles were evident for the newly characterised PnPro1, PnAda1, PnEbr1 and the carbon-catabolite repressor PnCreA. Overall, the results uphold PnPf2 as the central transcriptional regulator orchestrating genes that contribute to virulence on wheat and provide mechanistic insight into how this occurs.

**Importance:** Fungal pathogens cause large crop losses worldwide and consequently much attention has focused on improving host genetic resistance to diseases. These pathogens use effectors, which require coordinated expression at specific stages of the pathogenic lifecycle, to manipulate the host plant metabolism in favour of infection. However, our understanding of the underlying regulatory network in coordination with other genes involved in fungal pathogenicity is lacking. The Pf2 TF orthologues are key players underpinning virulence and effector gene expression in several fungal phytopathogens, including *P. nodorum*. This study provided significant insight into the DNA-binding regulatory mechanisms of *P. nodorum* PnPf2, as well as further evidence that it is central to the coordination of virulence. In the context of crop protection, the Pf2 taxonomic orthologues present opportune targets in major fungal pathogens that can be perturbed to reduce the impact of effector triggered-susceptibility and improve disease resistance.

## 1. Background

The *Parastagonospora nodorum*-wheat interaction has become a model fungal-plant pathosystem to study the molecular virulence factors underpinning infection. The fungus produces small secreted effector proteins that selectively interact with host-receptors encoded by dominant susceptibility-genes (1, 2). These interactions occur in a gene-for-gene manner that causes ‘effector-triggered susceptibility’ in the host plant, quantitatively affecting the disease which manifests as septoria nodorum blotch. Several effectors acting in this manner have now been identified and characterised for their role in virulence (3–7). These studies have also described a consistent pattern: the expression of these genes is maximal two to four days after infection and then declines. Furthermore, expression levels can vary by the presence or absence of their matching wheat receptors, as well as by epistasis, whereby one effector gene causes suppression of another (8–10). Yet relatively little is known concerning the mechanisms governing the effector gene regulation. In particular, are there common or distinct regulatory pathways involved? Do these components specifically control effector gene expression, or co-regulate other metabolic and developmental pathways? New knowledge in this area could present suitable targets to suppress for disease control.

Many fungi possess a Zn_2_Cys_6_ transcription factor (TF) Pf2 that has been associated with the regulation of effector gene expression. One example is the AbPf2 orthologue in *Alternaria brassicicola* that is critical for virulence on *Brassica* spp. (11). Gene deletion of *AbPf2* resulted in the down-regulation of a number of effector like genes, as well as putative cell-wall degrading enzymes. In *P*. *nodorum*, at least two key effector genes, *ToxA* and *Tox3*, require PnPf2 t be expressed (12). An RNA-seq analysis also revealed PnPf2 regulates many more putative effectors, CAZymes, hydrolases/peptidases and nutrient transporters (13). The PtrPf2 orthologue in *Pyrenophora tritici-repentis* controls *ToxA* expression and virulence on wheat, much like for the homologous *ToxA* gene in *P. nodorum* (12). In *Leptosphaeria maculans*, the causal agent of blackleg disease on *Brassica* spp., the LmPf2 orthologue also regulates several effector genes, including *AvrLm4-7, AvrLm6, AvrLm10A* and *AvrLm11*, as well as CAZyme expression (14).

The Pf2 orthologues can be traced across several Ascomycota fungal lineages that include the Dothideomycetes, Leotiomycetes and Soradiomycetes (15). Gene-deletion in the plant pathogens *Botrytis cinerea*, *Fusarium* spp., *Magnaporthe oryzae* and *Zymoseptoria tritici* all suppressed fungal virulence as well as their capacity to utilise alternative carbon sources (16–19). Analogous carbohydrate regulatory roles have been attributed in the saprophytic fungi *Neurospora crassa* and *Trichoderma reesei* (20, 21). In *N. crassa*, the putative orthologue Col-26 is a critical component within a signalling-network that responds to glucose availability and promotes the expression of CAZymes for plant cell-wall degradation (22–24). There is a strong association between CAZyme gene content and plant-pathogenic lifestyles (25).

There are some key factors yet to be established among Pf2 orthologues. Which DNA-regulatory elements are bound? Are Pf2-regulated genes directly targeted or is their expression modulated by indirect factors? Are there other TFs intimately connected with Pf2 regulation of virulence? The research presented here provides such critical insight for PnPf2 in *P. nodorum*, and establishes its direct role in effector expression and CAZyme regulation. A subsequent functional investigation of several connected and directly targeted TFs provides further evidence for the central role held by PnPf2 in the transcriptional network that orchestrates virulence in *P. nodorum*.

## 2. Results

### 2.1. PnPf2 possesses typical Zn_2_Cys_6_ TF domains and localises to the nucleus

Conserved domains or distinguishing features were analysed in the 652 amino acid (a.a) PnPf2 protein. The Zn_2_Cys_6_ DNA binding domain was located N-terminally at a.a 9 to 54 with an overlapping nuclear localisation signal (NLS) (KKGPKGSR; a.a 51 to 58) (**Fig. 1A**). A ‘fungal TF domain’ was identified from a.a 223 to 294 within a conserved ‘middle homology region’ (a.a 104 to 320). These features are frequently observed in Zn_2_Cys_6_ TFs and have been linked to the modulation of TF activity (26, 27, 15). A structurally disordered domain, typically associated with post-translational modifications and intermolecular interactions (28), was also identified at the C-terminus of PnPf2. Together, these features suggested that PnPf2 possesses the typical functional domains underpinning DNA-binding Zn_2_Cys_6_ TF activity (26). Nuclear localisation of the C-terminally tagged PnPf2-GFP fusion protein was also observed (**Fig. 1B**). Along with the functional domains identified, this supported DNA-binding regulatory activity by PnPf2.

**Fig. 1.**
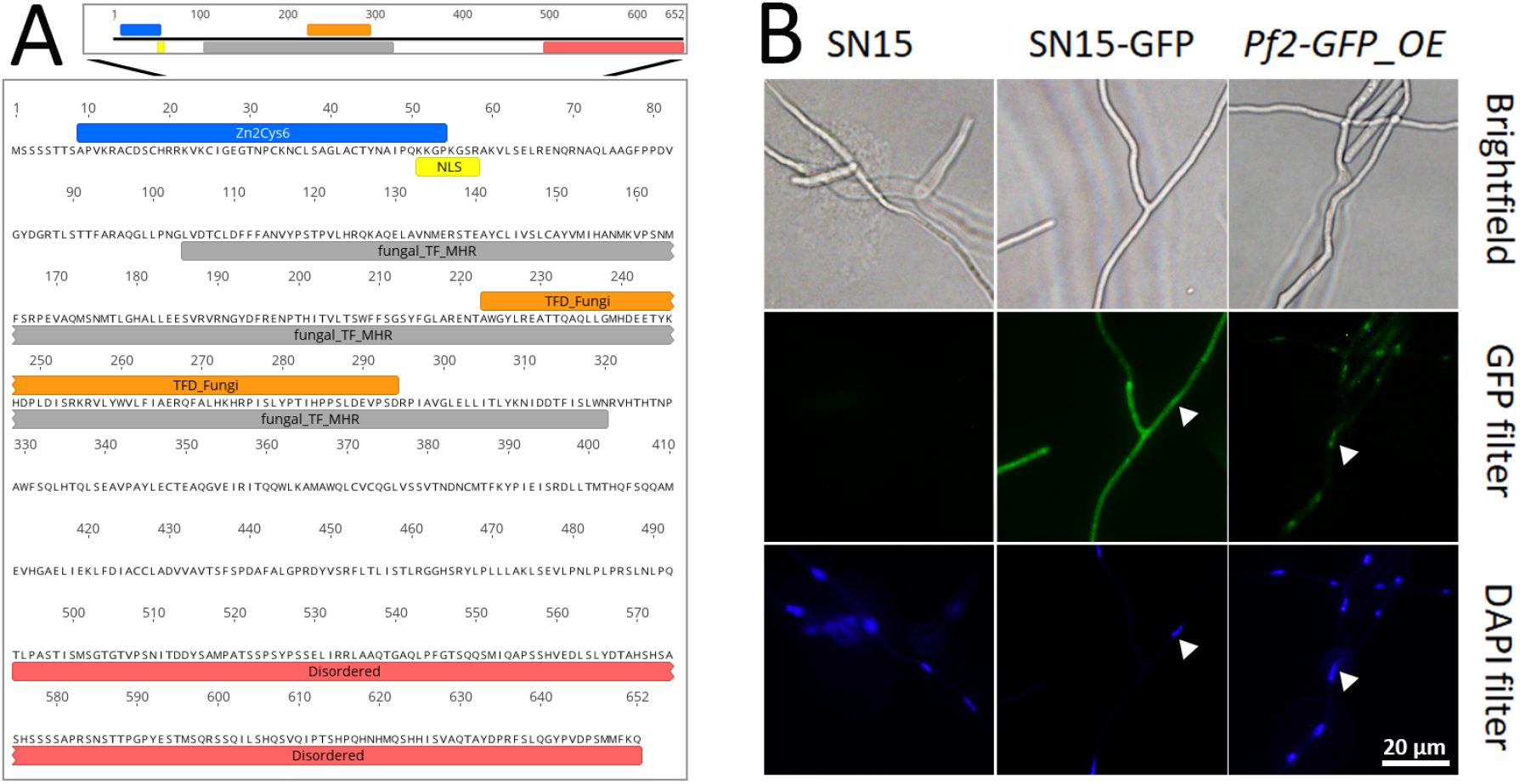
PnPf2 domain analysis and cellular localisation. A) Predicted domains and features identified in the 652 amino acid PnPf2 protein sequence typical of a Zn_2_Cys_6_ transcription factor (TF). The region corresponding to the N-terminal Zn_2_Cys_6_ DNA binding domain (Zn_2_Cys_6_ - Interpro IPR001138) is depicted in blue and the nuclear localisation signal (NLS) is in yellow. The fungal transcription factor domain (TFD_fungi - Interpro IPR007219) corresponds to orange within the ‘middle homology region’ (fungal_TF_MHR - Conserved Domain Database CD12148) depicted in grey. A C-terminal disordered protein region lacking secondary structure was identified that corresponds to the red bar. B) Epifluorescence microscopy depicting nuclear localisation of the GFP-tagged PnPf2 translational fusion specific to the *Pf2-GFP_OE* overexpression strain, in contrast to the wildtype (SN15) and the positive control strain expressing cytoplasmic GFP (SN15-*GFP*). Arrows indicate the corresponding locations of fungal nuclei under the respective filters determined by DAPI staining of a germinated pycnidiospore. A fluorescence signal was not detected in the *Pf2-GFP* strain, where expression was driven by the native *PnPf2* promoter, indicating PnPf2 accumulates at relatively low abundance.

### 2.2. Two direct PnPf2 target motifs are associated with gene-regulation

A chromatin immunoprecipitation (ChIP) analysis was used to define PnPf2-DNA binding *in situ*. The *Pf2-HA* (native promoter) and *Pf2-HA_OE* (overexpression promoter) strains expressing the 3x haemagglutinin (HA) tagged PnPf2-HA fusion protein retained PnPf2 function, in contrast to a *pf2-HA_KO* deletion control (**Text S1**). A ChIP-seq analysis was therefore undertaken using the *Pf2-HA* and *Pf2-HA_OE* strains to identify ‘summits’ within enriched ‘peak’ regions. Summits corresponded to the best estimate of PnPf2-DNA binding loci within the peaks (29). The *Pf2-HA* and *Pf2-HA_OE* samples provided complementary datasets; the overexpression mutant to compensate for lower *PnPf2* expression under culture conditions (13, 30) and broaden the DNA interactions captured. A total of 997 summits were obtained from the *Pf2-HA* dataset and 2196 from *Pf2-HA_OE*, corresponding to 740 and 1588 peak regions respectively. There were 588 shared peaks identified between the samples (**File S1**), indicating strong overlap between the predicted PnPf2-targeted regions. The *Pf2-HA_OE* dataset broadened the scope of putative binding sites. A quantitative PCR (qPCR) analysis was then undertaken on independently prepared ChIP DNA, comparing the *Pf2-HA* and *Pf2-HA_OE* samples to the *pf2-HA_KO* control. Fold-enrichment values relative to *pf2-HA_KO* strongly correlated with ChIP-seq summit - Log_10_(Q-values), a proxy measure for PnPf2-DNA binding affinity, in both the *Pf2-HA* (P < 0.01 with Pearson’s r = 0.77) and *Pf2-HA_OE* (P < 0.01 with Pearson’s r = 0.74) datasets (**Text S1**). The high reproducibility across separate methodologies provided confidence in the robustness of ChIP-seq summit calls.

**Text S1** Chromatin immunoprecipitation (ChIP) strain assessment and overview of ChIP-seq/ChIP-qPCR.

**File S1** A spreadsheet detailing the genomic coordinates for ChIP-seq peak regions [columns A-D], the respective summit loci [E], the pileup height of the mapped reads [F] and the summit -Log_10_(Q-values) representing the difference of ChIP reads relative to the input control sample [G]. The *Pf2-HA* strain encompasses columns A-G and the *Pf2-HA_OE* strain encompasses column H-N. Also included are the genomic coordinates for the peak regions obtained by merging the overlapping regions from the *Pf2-HA* and *Pf2-HA_OE* samples using MAnorm (65) [O-S].

Previous RNA-seq differential-expression analyses had identified an enriched consensus motif (5’-WMGGVCCGAA-3’) in the promoter regions of both AbPf2 and PnPf2-regulated genes (11, 13). Despite harbouring the typical ‘CGG’ Zn_2_Cys_6_ binding triplet (26), an interaction with PnPf2 was not observed in a heterologous system, indicating regulatory cofactors may be required (31, 13). A search for DNA-regulatory elements that interact with PnPf2 from the ChIP-seq dataset identified two enriched motifs in the merged *Pf2-HA* and *Pf2-HA_OE* peak regions (**Fig. 2A**). The first motif designated as M1 (5’-RWMGGVCCGA-3’) closely matches the consensus motif from AbPf2 and PnPf2-regulated gene promoters (11, 13). The second motif designated as M2 (5’-CGGCSBBWYYKCGGC-3’) is novel for PnPf2, encompassing two copies of the canonical ‘CGG’ Zn_2_Cys_6_ binding triplets (26), separated by eight nucleotides. Interestingly, M2 matches the AmyR regulatory response element that was modelled in *A. nidulans* (32). Both M1 and M2 are close to the ChIP-seq summits for the *Pf2-HA* and *Pf2-HA_OE* datasets (**Fig. 2B**), suggesting they accurately reflected DNA-binding loci.

**Fig. 2.**
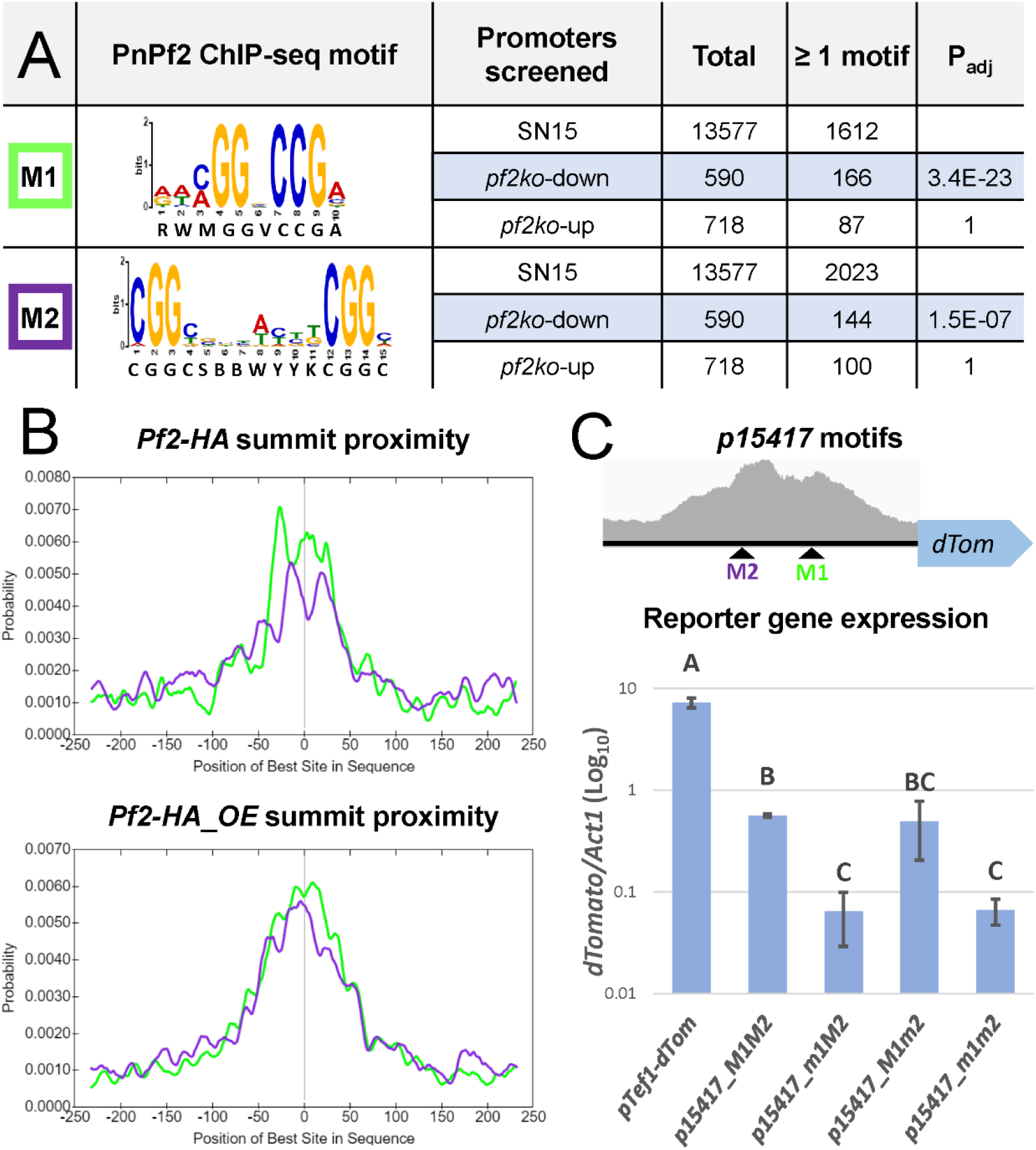
Identification of PnPf2 regulatory element motifs. **A)** The M1 motif (5’-RWMGGVCCGA-3’) and M2 motif (5’-CGGCSBBWYYKCGGC-3’) were modelled from the merged set of *Pf2-HA* and *Pf2-HA_OE* sample ChIP-seq peak regions. Their detection (≥ 1 occurrence) in the promoters of PnPf2 positively (*pf2ko*-down) or negatively (*pf2ko*-up) regulated gene promoters (13) are indicated relative to all SN15 promoters. The P_adj_ value reflects the test for significant enrichment (Fisher’s test with Bonferroni P_adj_ < 0.01), where both motifs were enriched in the *pf2ko*-down genes relative to SN15. **B)** The position of motif occurrences relative to ChIP-seq summits, demonstrating their higher likelihood at close proximity to the best estimate of PnPf2-DNA binding loci. **C)** Gene expression analysis assessing the effect of M1 and M2 motif mutation in *P. nodorum in situ*. The motif loci within a ChIP-seq peak in the SNOG_15417 gene promoter region (*p15417*) are depicted. The *dTomato* reporter gene was fused to a constitutive promoter control (*pTef1-dTom*) or the SNOG_15417 gene promoter (*p15417_M1M2*) in the *P. nodorum* background. The motifs were also mutated through substitution of the respective ‘CGG’ triplets, alone or in combination (*p15417_m1M2*, *p15417_M1m2* and *p15417_m1m2*). The strains *p15417_m1M2* and *p15417_m1m2* where M1 had been mutated exhibited significantly reduced expression relative to the non-mutated promoter in *p15417_M1M2*. This suggested PnPf2 regulatory activity had been impaired following M1 mutation but not M2 mutation in *p15417*. Letters indicate statistically distinct groupings by ANOVA with Tukey’s-HSD (P<0.05).

The previous RNA-seq analysis had defined genes positively or negatively-regulated by PnPf2 from their expression changes in the *PnPf2*-deletion mutant *pf2ko* relative to wildtype SN15 (13). These gene sets include the *in vitro* culture conditions that were replicated here for ChIP to maximise TF-DNA yields, as well as an early fungal infection stage *in planta* (72 hrs). Instances of M1 and M2 were then identified across the promoters of the positive (i.e. *pf2ko*-down) and negative (*pf2ko*-up) PnPf2-regulated gene sets (**File S2**). Both M1 and M2 were significantly enriched in the positively-regulated gene promoter set only (**Fig. 2A**). This indicates both motifs correspond to cis-regulatory elements that induce, rather than repress, gene expression.

A novel approach here sought to detect specifically the PnPf2-motif interactions. This utilised a *dTomato* reporter gene fused to the promoter of SNOG_15417. The promoter was chosen to encompass a ChIP-seq peak region with both the M1 and M2 motifs for a gene that is positively-regulated by PnPf2 (**Fig. 2C**; **File S2**). Integration of the construct at a predefined genomic locus in the SN15 background permitted evaluation of the reporter-gene expression in the resultant strain (*p15417_M1M2*) in comparison with strains where the CGG triplets in M1 and/or M2 had been substituted (*p15417_m1M2*, *p15417_M1m2* and *p15417_m1m2*). Significantly reduced expression was observed in the strains where M1 had been mutated, indicating that it is a functionally important and direct PnPf2 target in the SNOG_15417 promoter (**Fig. 2C**). No significant expression change was detected where only the M2 motif was mutated.

### 2.3. PnPf2 directly targets genes associated with the pathogenic lifestyle of *P. nodorum*

Genes with a ChIP-seq summit in their promoter region, considered a putative PnPf2 target, were cross-referenced with the *pf2ko* RNA-seq data analysis (**File S2**). There were 1286 targets identified from either ChIP-seq dataset, 484 of which were considered ‘high-confidence’ with a promoter summit in both *Pf2-HA* and *Pf2-HA_OE* (**Fig. 3A**). Of the 484 direct targets, 72 genes were positively-regulated in contrast to 6 negatively-regulated genes under the same *in vitro* conditions used for ChIP-seq. This indicates PnPf2 functions mainly as a positive regulator of gene expression. When expanded to also encompass PnPf2-regulated genes *in planta*, 93 were positively while 27 were negatively-regulated genes (**Fig. 3B**). Differential expression was not detected in the pf2ko mutant for 364 genes. This suggests other regulatory factors play a considerable role in their expression.

**File S2** A spreadsheet of PnPf2 regulation data across the *P*. *nodorum* SN15 genome for the respective annotated genes [column A]. Listed are whether ChIP-seq promoter summits were called from the *Pf2-HA* and *Pf2-HA_OE* samples [B-C], whether the enriched PnPf2 target motifs [D-E] or the putative PnCreA motif [G] were present in the gene promoter regions, and whether the gene was also down-regulated in the *pf2ko* mutant [G]. Also listed are the functional annotations [G-M]; whether the gene was classed as effector-like [H], a TF [I], the associated GO IDs/terms [J-K] and Interpro domain information [L-M]. The final columns list the respective gene expression data for *pf2ko* compared with SN15 either *in vitro* (*iv*) or *in planta* (*ip*) [13-22]. *Information indicated was derived from Jones et al. (2019) for comparative purposes.

**Fig. 3.**
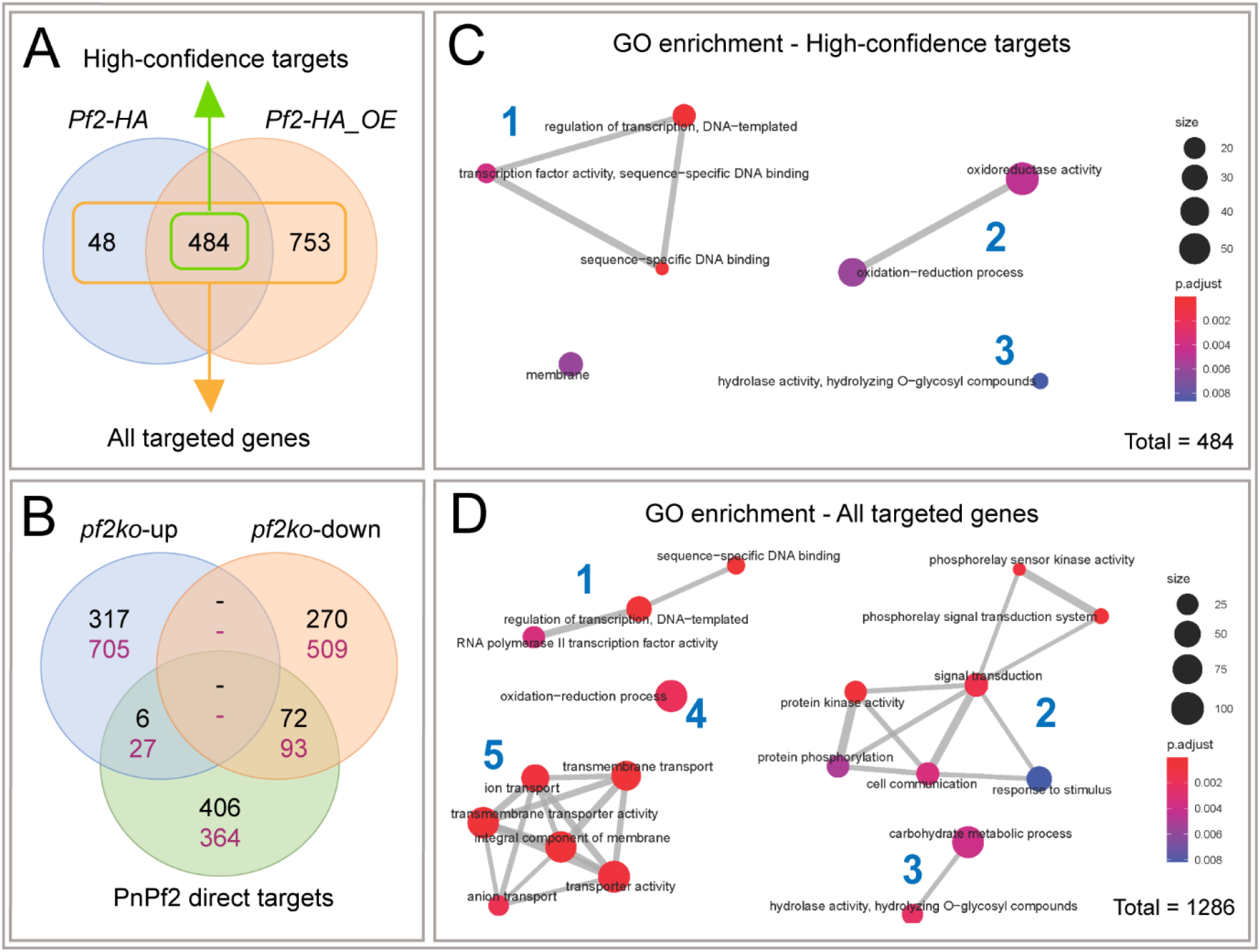
Gene expression and gene-ontology (GO) analysis of PnPf2 direct targets. **A)** Overview of the genes targeted by PnPf2 in their promoter region based on the respective *Pf2-HA* and *Pf2-HA_OE* ChIP-seq datasets. There were 484 genes considered high-confidence PnPf2 targets among the 1286 putative targets from either construct. **B)** The high-confidence targets in comparison with their expression pattern in the *pf2ko* deletion mutant. Black numbers correspond to the *in vitro* growth conditions replicated for ChIP-seq sample preparation while purple numbers also encompass differentially expressed genes during early infection. The greater overlap with PnPf2 positively-regulated (*pf2ko*-down - 72 genes) than repressed (*pf2ko*-up - 6 genes) suggested PnPf2 predominantly functions as a positive regulator of gene expression. **C-D)** A summary of significantly-enriched GO terms among PnPf2-targeted genes. The high-confidence (*Pf2-HA* and *Pf2-HA_OE*) and total identified targets (*Pf2-HA* or *Pf2-HA_OE*) are both displayed for comparison. Bubble sizes are proportionate to gene counts, colours to the enrichment test *P* values and the lines between bubbles to the total shared terms. Numbers in blue indicate connected gene networks representing transcription factors [number 1], redox molecules [2], carbohydrate-active enzymes [3], cell-signalling molecules [4], and trans-membrane transporters [5].

The characterised effector genes present in *P*. *nodorum* SN15, *ToxA, Tox1, Tox3* and *Tox267* (7), plus 29 other effector-like genes whose expression was altered in the *pf2ko* mutant, were assessed for evidence of direct regulation by PnPf2. Two distinct ChIP-seq summits were identified in the bi-directional *Tox3* promoter (**Fig. S1**). Both the upstream gene (i.e. SNOG_08982, encoding a protein disulphide-isomerase) and downstream gene (*Tox3*) are positively regulated by PnPf2. A ChIP-seq summit was also identified in the *Tox1* promoter, but only from the *Pf2-HA_OE* dataset (**Fig. S1**). Unlike Tox3, Tox1 necrosis-inducing activity is still detected in the *pf2ko* background (12), indicating the summit may represent a weak enhancer element. The *ToxA* gene is only expressed during infection but in a PnPf2-dependent manner. A weak promoter summit was observed despite multiple instances of the M1 motif, suggesting another factor(s) is required to facilitate PnPf2-DNA binding that was absent under the ChIP-seq experimental conditions. No distinct PnPf2 summit was observed in the promoter of *Tox267*, whose expression is not significantly altered in the *pf2ko* mutant, although two instances of M1 were identified >1000 bp upstream (**Fig. S1**). In total, 11 of the 29 PnPf2-regulated effector-like genes showed evidence of direct PnPf2-promoter binding through ChIP-seq summits (**Table S1-A**).

**Fig. S1** A depiction of the PnPf2 targeting of characterised effector genes in *P. nodorum* SN15. The *Pf2-HA* and *Pf2-HA_OE* ChIP-seq read peaks are presented at the *Tox3*, *Tox1*, *ToxA* and *Tox267* promoters. Peak summits were evident in the *Tox3* and *Tox1* promoters. Red dots represent instances of the M1 motif (5’- RWMGGVCCGA-3’) and blue dots M2 (5’-CGGCSBBWYYKCGGC-3’).

A gene-ontology (GO) enrichment and network analysis was then carried out to identify major functional gene classes that are directly regulated by PnPf2. Five distinct groups representing TFs, redox molecules, CAZymes, cell-signalling molecules and nutrient transporters were significantly enriched among the GO networks (**Fig. 3C-D**). The enrichment of CAZymes, redox molecules and nutrient transporters is consistent with enriched functional GO classes that were observed for the *pf2ko* differentially expressed genes (13). In contrast, the TFs and cell-signalling molecules were not enriched, indicating they could be tightly controlled by redundant pathways in addition to PnPf2. Nevertheless, it was striking that TF genes were particularly enriched in the high-confidence set of 484 targets (**Fig. 3C-D**). They made up 9.1% of these genes in contrast to 3.5% of the total genes annotated for SN15. Five TFs were directly targeted and positively regulated, providing a direct connection with PnPf2 in the regulation of virulence (**Table S1-B**).

### 2.4. PnPf2 is the central transcriptional regulator of host-specific virulence

The identification of TFs as major PnPf2 targets prompted a functional exploration of other TFs with a putative intermediate role in regulating virulence. Three directly-regulated TF genes were therefore targeted for deletion where orthologues had virulence-associated roles (**Table 1**). These included SNOG_03490 (*PnPro1*), SNOG_04486 (*PnAda1*) and SNOG_08237. Two additional TF-encoding genes were simultaneously investigated. This included SNOG_08565, identified in the PnPf2 lineage of fungal TFs (15) suggesting a possible common evolutionary origin. It also included SNOG_03067 (*PnEbr1*), which is co-expressed with *PnPf2, ToxA*, *Tox1* and *Tox3* high during early infection (**Fig. S2**), suggesting a similarly important role in disease.

**Table 1.** Rationale for the investigation of novel transcription factors (TFs) in this study

| TF investigated | Involvement | Virulence-associated orthologues <sup>A</sup> |
| --- | --- | --- |
| *PnPro1*<br>(SNOG_3490) | Directly-positively regulated by PnPf2 | *AbPro1* (*Ab*), *MoPRO1* (*Mo*), *GzZC232* (*Fg*), *UvPro1* (*Uv*) |
| *PnAda1*<br>(SNOG_04486) | Directly-positively regulated by PnPf2 | *GzbZIP001* (*Fg*), *FpAda1* (*Fp*) |
| SNOG_08237 | Directly-positively regulated by PnPf2 | *CoHox1* (*Co*) |
| SNOG_08565 | Shared ancestral lineage with PnPf2 orthologues | - |
| *PnEbr1*<br>(SNOG_03067) | Co-expressed with *PnPf2*, *ToxA*, *Tox1* and *Tox3* during infection | *EBR1* (*Fg*), *EBR1* (*Fo*), *MoCod2* and *Cnf2* (*Mo*) |
| *PnCreA*<br>(SNOG_13619) | Enriched CreA-binding motif (22) in PnPf2-regulated gene promoters | *CreA* (*Af*), *Cre1* (*Fo*), *CreA* (*Pe*) |

<sup>A</sup> Putative orthologues were inferred by cross-referencing a previous TF-orthology analysis and literature review (15, 36). Abbreviations: *Ab*; *Alternaria brassicicola*, *Af*; *Aspergillus flavus*, *Co*; *Colletotrichum orbiculare*, *Fg*; *Fusarium graminearum*, *Fo*; *Fusarium oxysporum*, *Fp*; *Fusarium pseudograminearum*, *Mo*; *Magnaporthe oryzae*, *Pe*; *Penicillium expansum*.

**Fig. S2** A heatmap depiction of *Parastagonospora nodorum* SN15 hierarchical cluster analysis. Clustering was based on microarray gene-expression data during infection (*in planta*) or axenic (*in vitro*) growth obtained from a previous study (30). Clusters were cut into the 10 most distant clusters to identify genes co-expressed with *PnPf2*, *ToxA*, *Tox1* and *Tox3*, which included the Zn_2_Cys_6_ transcription factor *PnEbr1* (SNOG_03037) therefore investigated in this study.

Gene deletion strains for the five TFs were phenotypically characterised in comparison to wildtype SN15 and *pf2ko*. The *pro1_KO*, *ada1_KO* and *ebr1_KO* deletion mutants presented distinct phenotypic abnormalities (**Fig. 4**). The *pro1_KO* mutants were abolished in their ability to form pycnidia and sporulate both during infection and on nutrient-rich agar. However, vegetative growth was expansive in both conditions (**Fig. 4A-B**), suggesting PnPro1 acts to suppress hyphal development. Although *PnPro1* is positively-regulated by PnPf2, there was no distinct phenotypic overlap with the *pf2ko* mutant. The *ada1_KO* mutant was significantly reduced in virulence on detached leaves (**Fig. 4A-B**). Dark brown discolouration at the site of infection suggested a hypersensitive response had contained the infection. We also observed an increased susceptibility to oxidative (H_2_O_2_) stress for *ada1_KO* mutants similar to *pf2ko*. Furthermore, sporulation was reduced in *ada1_KO* relative to SN15 (**Fig. 4C-E**). The *ebr1_KO* mutants exhibited vegetative growth defects with an uneven growth perimeter around the colony edges coinciding with perturbed virulence (**Fig. 4A-B**). Similar hyphal-branching defects were described following deletion of *PnEbr1* orthologues in *Fusarium* spp. (33, 34). Interestingly, the *ebr1_KO* mutants were also susceptible to H_2_O_2_ stress at a level comparable to *pf2ko* and *ada1_KO*. Furthermore, pycnidia were abnormally developed, although still viable for the production of conidia, but were not detected on infected leaves (**Fig. 4C-D**). We did not observe morphological or virulence defects for the *08237_KO* or *08565_KO* mutants (**Text S2**) which were not investigated in further detail.

**Fig. 4.**
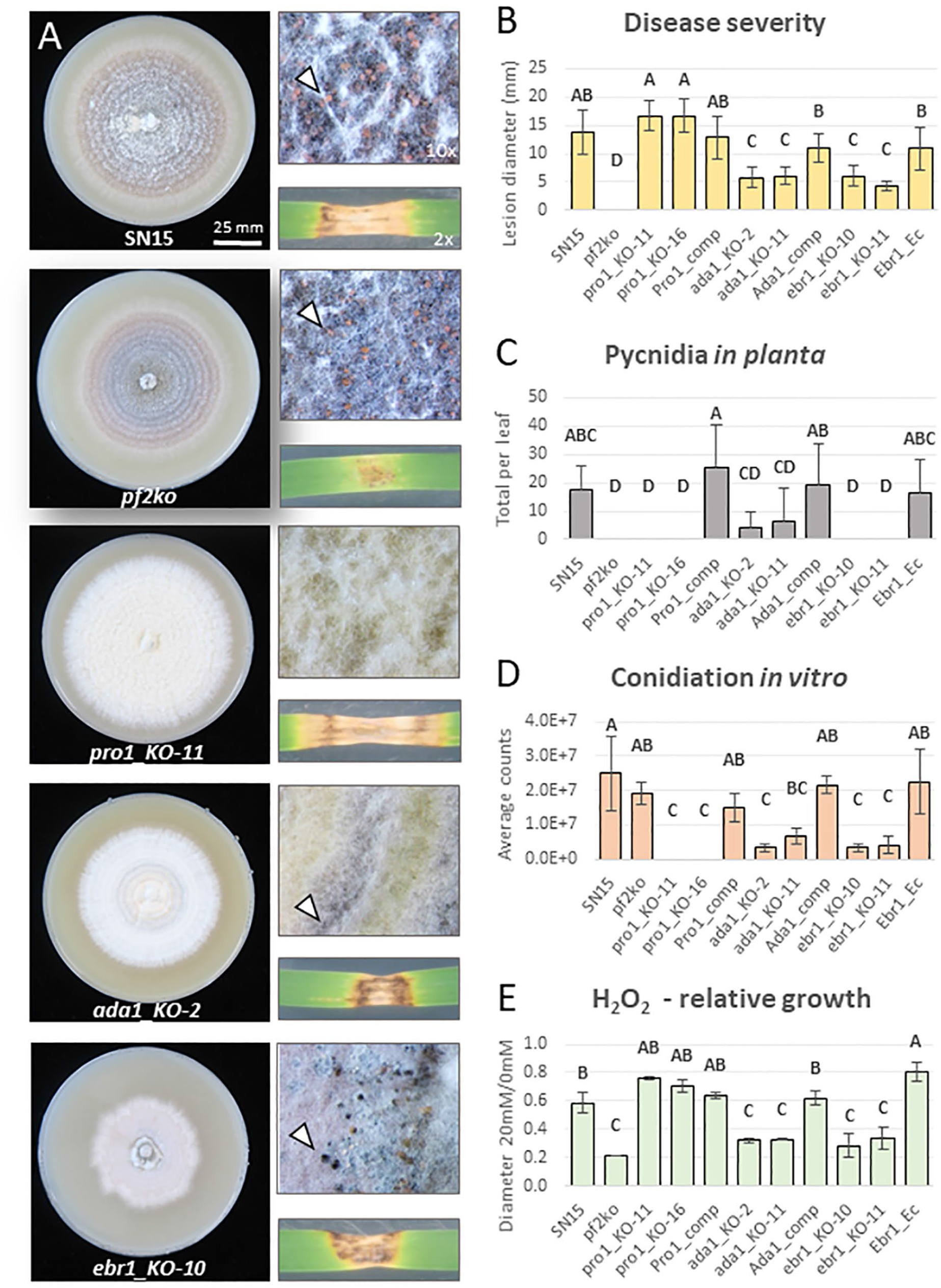
Phenotypic assessment of transcription factor (TF) gene deletion mutants. **A)** Representative images after 12 days of growth on nutrient-rich agar (V8PDA) and infection on detached wheat leaves (cv. Halberd). Arrows demonstrate pycnidia if they were detected in the respective mutants. **B)** Average lesion sizes representing disease severity. **C)** Pycnidia counts, a measure of pathogenic fitness following the infection. **D)** Average conidial (pycnidiospore) counts on V8PDA. **E)** Growth inhibition on 20mM H_2_O_2_ relative to 0mM on minimal medium agar. Letters indicate statistically distinct groupings by ANOVA with Tukey’s-HSD (P<0.05).

During the course of this study the carbon-catabolite repressor (CCR) element was modelled as the binding site for the Cre-1 TF that suppresses CAZyme expression in *N. crassa* (22). We noted this was near identical to a motif (5‘-RTSYGGGGWA-3’) that is also enriched in PnPf2-regulated gene promoters (13) but not identified from the ChIP-seq peaks. Since Cre-1 orthologues are conserved CCR regulators in filamentous fungi (35, 36), and since the CCR element is also enriched in PnPf2 regulated gene promoters, a putative Cre-1 orthologue (PnCreA) was investigated in *P. nodorum* to identify common regulatory pathways with PnPf2. Both *PnCreA* overexpression and gene-deletion mutants (*CreA_OE* and *creA_KO*) were created and then investigated alongside *pf2ko* and a *PnPf2* overexpression mutant (*Pf2_OE*). Despite clear phenotypic-growth abnormalities (**Fig. 5**), neither the *CreA_OE* nor *creA_KO* mutants exhibited virulence defects on wheat leaves (**Text S2**). The *creA_KO* strain was enhanced in starch utilisation (**Fig. 5**), an indicator substrate for CCR activity (37). In contrast, there was a moderate reduction of *pf2ko* to utilise starch, similar to observations in other fungal *PnPf2*-orthologue mutants (17, 19, 21). These results support contrasting roles between PnCreA and PnPf2 for the regulation of some CAZyme-related genes.

**Text S2** Supplemental transcription factor mutant phenotype assessment.

**Fig. 5.**
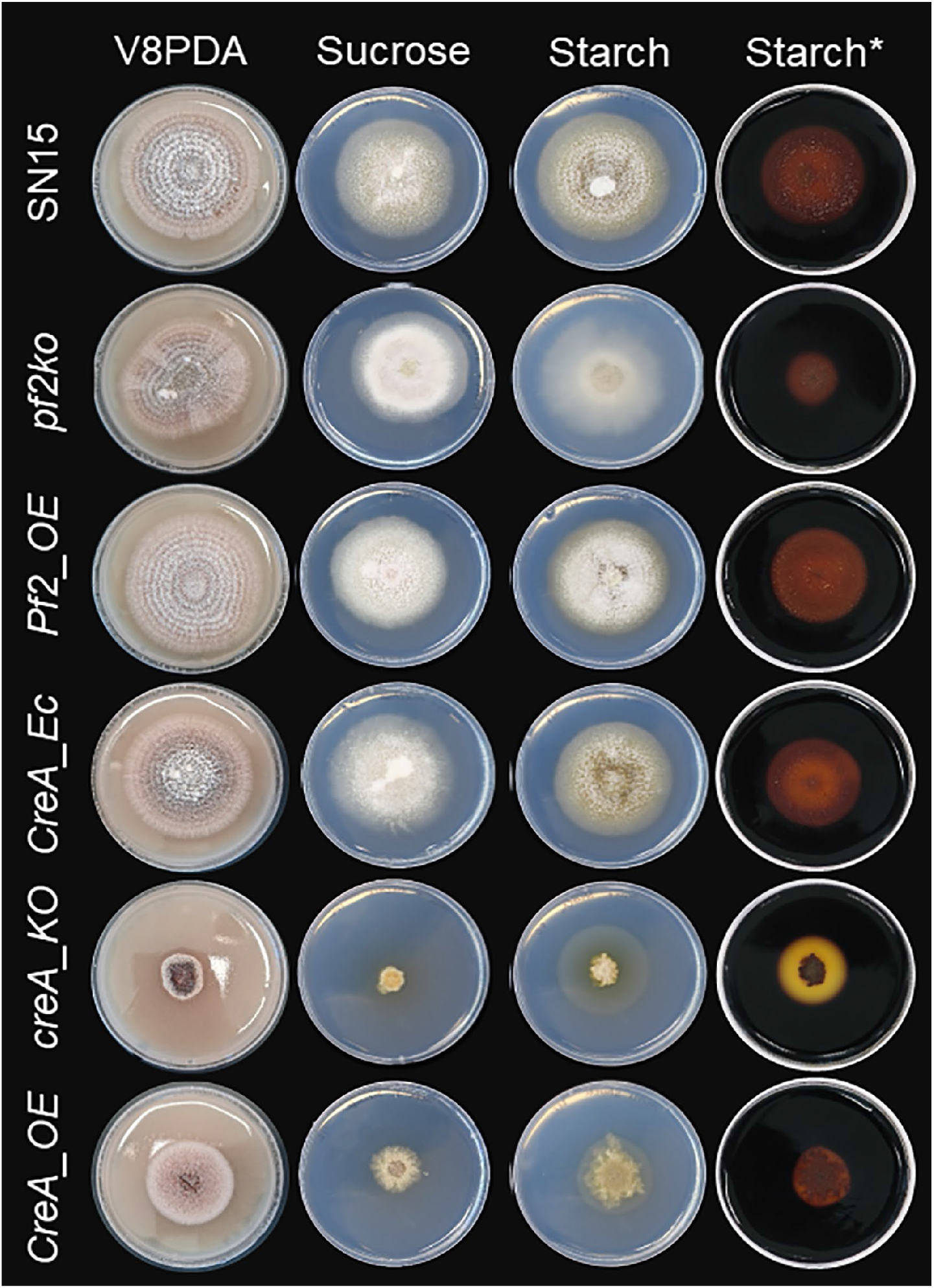
Assessment of *PnPf2* and *PnCreA* mutant growth on different substrates. Images following 12 days of growth on nutrient-rich V8PDA and minimal-medium agar with a primary (sucrose) or secondary (starch) carbon source. Wildtype SN15 and the respective mutants are listed on each row. The capacity to fully utilise starch was enhanced in the *creA_KO* mutant despite growth defects, suggesting impaired carbon-catabolite repression. Starch utilisation was moderately reduced in the *pf2ko* mutant. *Post-stained with Lugol’s iodine to assist visualisation of starch hydrolysis.

## 3. Discussion

Prior to this research, PnPf2 had been identified as an important regulator of *P. nodorum* virulence on wheat (12, 13), but important details and mechanistic insights were missing. We sought to take further steps and establish the DNA-binding elements targeted by PnPf2 and identify genes that were directly under its direct regulation. Two distinct regulatory motifs M1 and M2 were identified and linked to positive gene-regulation by PnPf2. M1 was strikingly similar to an enriched sequence in AbPf2 positively-regulated gene promoters (11), possibly representing a conserved Pf2 binding mechanism. It will be pertinent to explore this motif as a regulatory target for other fungal Pf2 orthologues (11–14, 16–21). Interestingly, the M2 motif matches the extensively characterised AmyR regulatory-response element in *A. nidulans* (38, 32). Polysaccharide metabolism has long been established as a regulatory function for AmyR (39, 40). Therefore, some shared regulatory pathways likely exist with Pf2 orthologues given the evidence for at least one conserved binding mechanism. However, there are major a.a polymorphisms between AmyR and Pf2 orthologues at the Zn_2_Cys_6_ DNA-binding domain (15) and M1 has not been reported as an AmyR target despite extensive motif investigation (38, 32). It is therefore conceivable that M1 is a regulatory binding element unique to Pf2 orthologues and therefore useful to identify putative direct targets such as *ToxA* in *P. nodorum*.

The ChIP-seq PnPf2-DNA binding dataset also facilitated the identification of *P. nodorum* genes under direct PnPf2 regulation. Among these genes is the *Tox3* effector and the adjacent gene, SNOG_08982, encoding a protein disulphide isomerase. This class of protein catalyses cysteine-cysteine bond formation which has been connected to fungal effector protein production (41). Therefore, it would be worth exploring any involvement of SNOG_08982 in the post-translational modification of Tox3 and other effectors. PnPf2 binding was also detected in the *Tox1* promoter. A partial reduction in *Tox1* expression was reported in the *pf2ko* mutant (13), indicating PnPf2 is not essential but enhances expression under favourable conditions. *ToxA* is only expressed *in planta,* but is PnPf2 dependent (12). Despite multiple instances matching the M1 motif, there was little evidence for PnPf2-*ToxA* promoter binding, suggesting chromatin inaccessibility or the absence of essential binding-cofactors under the ChIP culture conditions. Direct PnPf2 regulation of *Tox267* was not evident. The other recently-cloned effector gene *Tox5* is not present in the SN15 isolate used in this study, but is homologous to *Tox3* may be under PnPf2 control (6). Nevertheless, several other effector-like genes were identified as direct PnPf2 targets (**Table S1-A**). Importantly this analysis provided strong evidence that PnPf2 is a key direct-regulator of effectors, the major *P. nodorum* virulence factors in the lifestyle of this pathogen.

Evidence for regulation of effector expression has been reported for another *P*. *nodorum* TF PnCon7 (42), yet its apparent requirement for fungal viability renders it difficult to investigate a precise functional role. Here, several novel TFs were functionally investigated based on their connection to PnPf2 (**Table 1**). We did not observe any change in the necrosis-inducing activity on wheat of fungal culture filtrates derived from the respective mutants. However, developmental virulence roles, including oxidative stress tolerance and hyphal development, were identified for *P. nodorum* PnAda1 and PnEbr1. It is possible that the direct regulation of PnAda1 by PnPf2 contributes to the susceptibility to oxidative stress also identified in the *pf2ko* mutant. The PnCreA orthologue of *N*. *crassa* Cre-1 was also investigated, following the striking observation that the *N*. *crassa* Cre-1 CCR element (5′-TSYGGGG-3’) was enriched in PnPf2-regulated gene promoters (13). Furthermore, Cre-1 and the PnPf2 orthologue Col-26 are both key components of a transcriptional network controlling CAZyme production in *N. crassa* (22–24, 43). Here, the *creA_KO* strain displayed an enhanced capacity to utilise starch, which was moderately impaired in the *pf2ko* mutant (**Fig. 5**). This indicates PnCreA and PnPf2 shared a similar function to the respective *N. crassa* orthologues (21). Surprisingly however, despite vegetative growth abnormalities on agar, there was no distinct change in the virulence profile of either the *creA_KO* or *CreA_OE* mutants (**Text S2**). We also failed to detect the CCR element in the promoters of *ToxA*, *Tox1*, *Tox3* or *Tox267* (**File S2**). This suggests that the regulation of host-specific virulence factors critical for *P. nodorum* infection are not subject to CCR by PnCreA.

This investigation, along with previous TF studies in *P. nodorum* (44, 45, 12, 42), indicate PnPf2 is central to the transcriptional-regulatory network controlling virulence, for which a tentative model is proposed (**Fig. 6**). Having expanded our understanding, it also raised some key questions. For many genes directly targeted by PnPf2, differential expression in *pf2ko* has not been observed (364 of 484 high-confidence targets). Such discrepancies are also reported in ChIP-seq experiments on filamentous fungi (**Table S2**). One aspect to consider is that functional TF binding requires specific cofactors/coregulators before gene expression is eventually modulated (46, 47). Furthermore, TF-DNA interactions can be redundant or non-functional (48–50). It is therefore plausible that many binding sites are transiently occupied by PnPf2 in this manner, acting as a biological sink. A change in the epigenetic landscape, for example during growth *in planta*, could open up genomic regions for which PnPf2 exhibits a high affinity and then actively binds. Performing PnPf2 ChIP during early infection will likely prove highly useful in this regard if sufficient fungal material can be obtained. ChIP-seq targeting histone marks specific for euchromatin or heterochromatin under infection conditions, or methylation-sensitive sequencing are alternatives to provide insight into the genome accessibility of PnPf2 (14, 51–53). The identification of both the M1 and M2 motifs carrying alternatively oriented ‘CGG’ triplets, typical of Zn_2_Cys_6_ monomers (26), was suggestive of PnPf2 dimerisation with other Zn_2_Cys_6_ TFs. However, deletion of the putative ancestral PnPf2 homologue SNOG_08565 did not provide any phenotypic response that would indicate a connection. Therefore, future investigations will do well to explore these interactions, for example through co-immunoprecipitation/affinity purification analysis or a yeast-2-hybrid screen, to delineate PnPf2-DNA binding mechanisms. Functional investigation of the PnPf2 ‘middle homology region’ and C-terminal disordered region may also provide insight into the upstream signalling pathways that activate or repress PnPf2 activity through these domains.

**Fig. 6.**
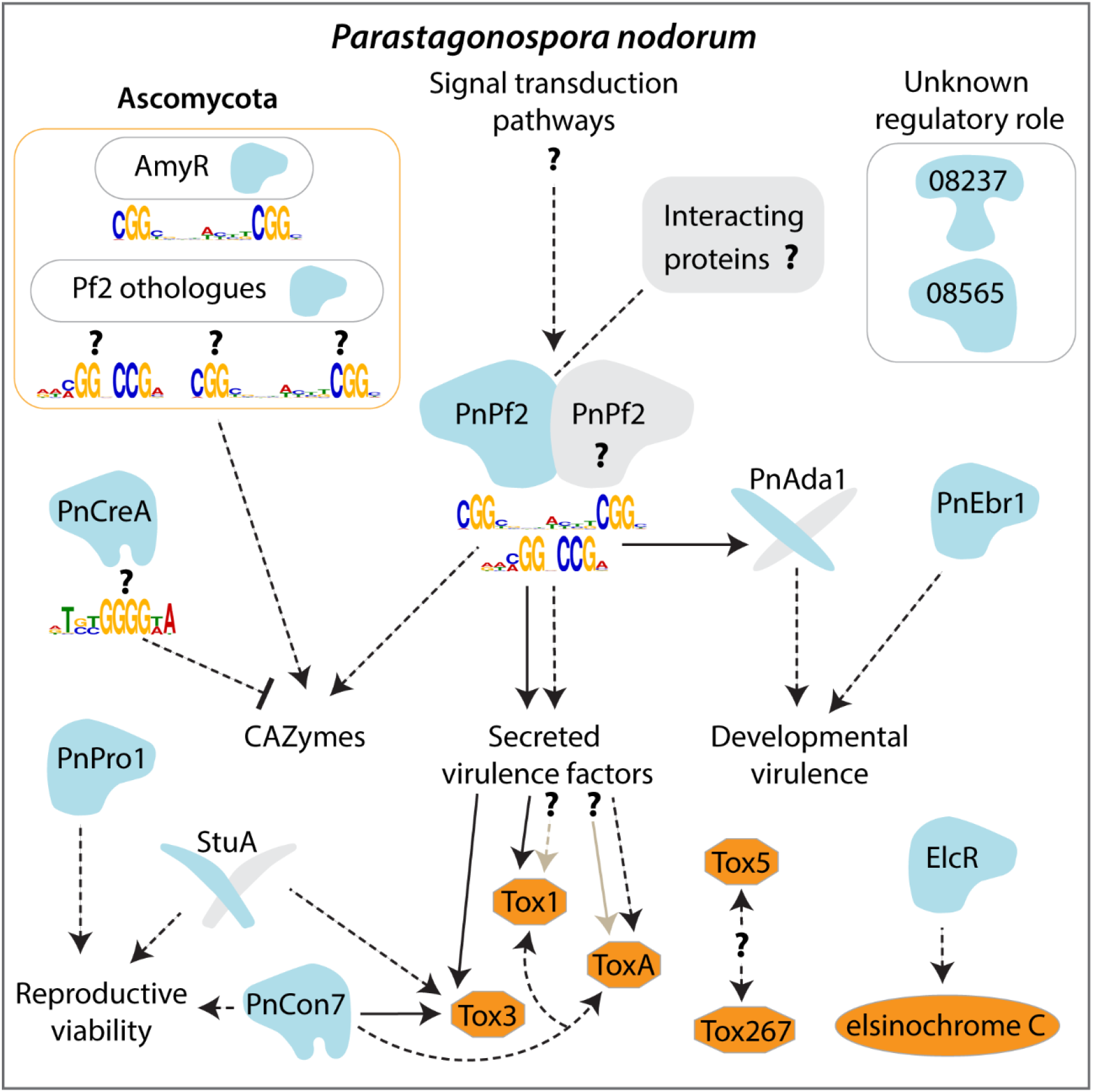
The proposed model of the PnPf2-centred regulatory network in the virulence of *P. nodorum*. The Pf2 taxonomic orthologues and AmyR regulators in the Ascomycota fungi (discussed in the main text) are presented for context regarding the two PnPf2 regulatory motifs described in this study. Transcription factors (TFs) and virulence factors that are cloned and characterised in *P. nodorum* are depicted (blue and orange shapes respectively; grey shapes indicate putative interacting proteins). Dashed arrows show where gene regulation occurs and solid arrows represent direct regulation. Question marks are presented where evidence is tentative and requires validation. PnPf2 controls the expression of key virulence factors. The effector *Tox3* is directly regulated and *ToxA*, based on promoter-motif and gene expression data, is likely a direct target during plant infection. PnPf2 also directly targets the *Tox1* promoter as a possible enhancer while regulators of *Tox267* and *Tox5* require investigation. Carbohydrate-active enzymes (CAZymes) are also regulated by PnPf2, with a subset putatively repressed by PnCreA for which no distinct role in virulence has been established. Developmental virulence, such as oxidative stress tolerance and hyphal growth, were processes attributed in this study to the PnPf2 targets PnAda1 and PnEbr1. PnPro1 and StuA (44) are essential for reproduction by sporulation, while no distinct role was identified for the putative TFs encoded by SNOG_08237 and SNOG_08565. Elsewhere, PnCon7 has been reported to regulate effector expression but is an essential viability factor, while production of a phytotoxic metabolite elsinochrome C is controlled by the pathway-specific ElcR gene-cluster TF (45, 42).

To conclude, this study presents direct evidence of DNA binding in a Pf2 orthologue, where virulence-regulatory functions are consistently observed in phytopathogenic fungi. In *P. nodorum*, PnPf2 is central to the transcriptional regulation of virulence and directly controls effector expression. The current research on PnPf2 now provides a platform to further investigate its signalling pathways and molecular interactions that could be inhibited for targeted disease control.

## 4. Materials and methods

### 4.1. Annotations and PnPf2 domain analysis

The *P. nodorum* annotated genome for the reference isolate SN15 (54) was used consistent with annotations in the previous RNA-seq analysis (13). The PnPf2 polypeptide sequence was submitted to Interproscan (Release 82.0) for Interpro and Conserved-Domain-Database domain identification (55). NLStradamus was used to predict the nuclear localisation signal (56). The disordered region was predicted using IUPRED2A (28).

### 4.2. Generation and assessment of fungal mutants

The molecular cloning stages, the constructs generated and diagrammatic overview of the final transformed *P*. *nodorum* mutants generated in this study are detailed in **Text S3**. The respective primers designed for fragment amplification and/or screening are outlined in **File S3**. Fungal mutants used in this study are summarised in **Table 2**. Their phenotypic and gene-expression analysis procedures are also described in **Text S3.**

**Table 2.** Overview of the strains referenced in this study ^A^

| Strain ID | Description <sup>B</sup> |
| --- | --- |
| SN15 | Wildtype *P. nodorum* reference isolate |
| *pf2ko* | Original *PnPf2* deletion mutant<br>(12, 13) |
| *pf2_KO* | *PnPf2* deletion mutant to facilitate targeted complementation |
| *Pf2_OE* | *PnPf2* overexpression (*pGpdA* promoter) |
| *Pf2-GFP* | *PnPf2* with *GFP* tag, native promoter |
| *Pf2-GFP_OE* | *PnPf2* with *GFP* tag, overexpression promoter |
| *Pf2-HA* | *PnPf2* with 3xHA tag, native promoter |
| *Pf2-HA_OE* | *PnPf2* with 3xHA tag, overexpression promoter |
| *pf2-HA_KO* | *PnPf2* coding sequence replaced with 3xHA tag |
| *CreA_OE* | *PnCreA* overexpression (*pGpdA* promoter) |
| *creA_KO* | *PnCreA* deletion mutant |
| *CreA_Ec* | *PnCreA* ectopically-integrated construct |
| SN15-GFP | SN15 constitutively expressing *GFP* |
| *pTef1-dTom* | SN15 constitutively expressing *dTomato* (defined locus) |
| *p15417_M1M2* | SN15 expressing *dTomato* (defined locus) SNOG_15417 promoter, no mutations |
| *p15417_m1M2* | SN15 expressing *dTomato* (defined locus) SNOG_15417 promoter, M1 mutated |
| *p15417_M1m2* | SN15 expressing *dTomato* (defined locus) SNOG_15417 promoter, M2 mutated |
| *p15417_m1m2* | SN15 expressing *dTomato* (defined locus) SNOG_15417 promoter, M1+2 mutated |
| *pro1_KO* | *PnPro1* deletion mutant |
| *Pro1_comp* | *PnPro1* complemented in *pro1_KO* background |
| *ada1_KO* | *PnAda1* deletion mutant |
| *Ada1_comp* | *PnAda1* complemented in *ada1_KO* background |
| *08237_KO* | SNOG_08237 deletion mutant |
| *08237_comp* | SNOG_08237 complemented in *08237_KO* background |
| *08565_KO* | SNOG_08565 deletion mutant |
| *08565_comp* | SNOG_08565 complemented in *08565_KO* background |
| *ebr1_KO* | *PnEbr1* deletion mutant |
| *Ebr1_Ec* | *PnEbr1* ectopically-integrated construct |

<sup>A</sup> Strains are listed in corresponding order to the detailed description of their generation in Text S3.
<sup>B</sup> Abbreviations: GFP – Green fluorescent protein, HA – Haemagglutinin, M1 – Motif1, M2 – Motif 2.

**Text S3** Supplemental materials and methods.

**File S3** A spreadsheet compilation of the primers used in this study organised by their general use [column 1] with the primer ID and sequence [2-3] and descriptions for their use [4-5]. Highlighted in italics are restriction enzyme recognition sites, in bold are overlapping regions used in cloning and in red are sites for incorporating single nucleotide changes during cloning.

### 4.3. Chromatin immunoprecipitation sample preparation

The *Pf2-HA*, *Pf2-HA_OE* and *pf2-HA_KO* strains were prepared following 3 days standardised growth in 100 mL Fries3 liquid medium (**Text S3**). Prior to harvesting, a 5 mL crosslinking solution (10% w/v formaldehyde, 20 mM EDTA and 2 mM PMSF dissolved in 50 mM NaOH) was added with continuous shaking at 100 rpm for 10 min. To this, 5 mL quenching solution (1.25 M glycine) was added before another 10 min shaking. Whole protein extracts were then obtained as described (**Text S3**) with modifications for ChIP. The 50 mM Tris was replaced with 50 mM HEPES in the lysis buffer while gentle rotation of the resuspended fungal material was replaced by eight rounds of sonication using a Bandelin (Berlin, Germany) UW3100+SH70+MS73 tip sonicator to fragment the fungal DNA (set at 15 sec on/off with 60% amp and 0.8 duty cycle). Samples were held in an ice block during sonication. The supernatant was then retrieved from two rounds of centrifugation (5000 g, 4 °C for 5 min). A 100 μL aliquot of the supernatant was reserved as an ‘input control’ against which ChIP samples were to be normalised. A 1000 μL aliquot was then precleared for immunoprecipitation by gently rotating with 20 μL Protein A dynabeads (10001D - Thermofisher, Waltham, Massachusetts) for 1 hr at 4 °C. The supernatant was then retrieved and incubated with 2.5 μg anti-HA polyclonal antibody (71-5500 - Thermofisher) for 16 hrs at 4 °C. Another 20 μL Protein A dynabeads were then added and gently rotated for 2 hrs at 4 °C. The dynabeads were then retrieved and washed twice with 1mL ice-cold lysis buffer, once with high-salt buffer (lysis buffer + 500 mM NaCl), once with LiCl buffer (250 mM LiCl, 10 mM Tris-HCl, 1 mM EDTA, 0.5% NP40 and 0.5% NaDOC) and once with TE buffer (10 mM Tris-HCl, 1 mM EDTA, pH 8). Samples were then incubated in a shaking incubator for 10 min (300 rpm, 65 °C) with 200 μL elution buffer (0.1 M NaHCO_3_, 10 mM EDTA and 1% SDS) before transferring the supernatant to a fresh tube. The input control was also supplemented with 100μL elution buffer at this stage and 8 μL NaCl solution (5 M) was added to both samples before de-crosslinking for 16 hrs at 65 °C. To these samples, 200 μL of H_2_O and 100 μg RNAse A (QIAGEN, Hilden, Germany) were added before incubating for 1 hr at 65 °C. Ten μg Proteinase K (Sigma-Aldrich, St. Louis, Missouri) was then added before incubating a further 1 hr at 50 °C.

For ChIP-qPCR, DNA (for both the *Pf2-HA*, *Pf2-HA_OE* and *pf2-HA_KO* ChIP and input control samples) was recovered from Proteinase K treated samples using the GenElute PCR purification kit (Sigma-Aldrich).

For ChIP-seq analysis, DNA (for both *Pf2-HA* and *Pf2-HA_OE* ChIP and input control samples) was purified from the Proteinase K treated samples by mixing in 1 volume (400 μL) of phenol:chloroform. This was centrifuged for 5 min at 16000 g and the aqueous phase retrieved. To this, 400μL chloroform was added, mixed and spun (16000 g 5 min) before 350 μL of the aqueous phase was transferred to a fresh tube. 35 μL sodium acetate (3 M, pH 5.2) was added with 1 μL of glycogen (20mg/mL). Samples were mixed by inversion and 1 mL 100% ethanol added before precipitation at −80 °C for 1-2 hrs. Pellets were retrieved by spinning 16000 g for 10 min at 4 °C, then washed in 1 ml of ice-cold 70% ethanol before drying and resuspension in 30 μL Tris-Cl (10 mM).

Two independent DNA preparations for each sample (i.e. the ChIP and input samples for both *Pf2-HA* and *Pf2-HA_OE*), beginning with the fungal growth stage in Fries3 broth, were pooled to ensure sufficient DNA was obtained for generating ChIP-seq libraries. The pooled DNA was measured using a Tapestation system (Agilent, Santa Clara, California). 10 ng of each sample was processed using the TruSeq ChIP Library Preparation Kit (Illumina, San Diego, California). Libraries were size-selected (100-300 bp) and split across four separate lanes for sequencing in a NextSeq 500 sequencer (Illumina) to obtain 2 x 75 bp paired-end reads (Australian Genome Research Facility, Melbourne, Australia).

### 4.4. ChIP-seq analysis

An overview of the following data analysis pipeline from QC of raw reads through to genome mapping, ChIP-seq peak/summit calling, target gene prediction, ChIP-qPCR validation, GO enrichment analysis and motif position-weight-matrix (PWM) modelling **Text S1**.

#### 4.4.1. Raw read filtering, mapping and peak/summit calling

Raw reads were checked using FASTQC (Version 0.11.9) (57) and the adapter sequences were trimmed using Cutadapt (Version 1.15) along with nucleotides where the Illumina quality scores were below 30 (58). Optical duplicates were then removed using the ‘dedupe’ option in Clumpify (version 1.15) from the BBTools package (59). Reads were subsequently mapped to the SN15 genome (54) using BWA-MEM (60). Reads mapping to a single locus as the best match (primary alignments) were retained for downstream analysis and the datasets from sample libraries originally split across the NextSeq lanes were merged using SAMtools (Version 1.10) to produce the final mapped-read datasets (61). MACS (Version 2.2.7.1) was used for calling enriched regions (i.e. peaks) and summits (highest nucleotide point or points within peak regions) from ChIP sample reads relative to the input samples (for *Pf2-HA* and *Pf2-HA_OE*). A Q-value peak enrichment threshold of 0.01 was used and the BAMPE option utilised to assess read depth from cognate pairs (62, 63). Paired read lengths from the cognate pairs were assessed using Deeptools ‘bamPEFragmentSize’ (Version 3.3.0) to verify they corresponded to 100-300 bp size selected fragments (64).

#### 4.4.2. Modelling binding-site motifs

The overlapping peak regions identified from the *Pf2-HA* and *Pf2-HA_OE* samples were merged using MAnorm (65) to create a consensus set of enriched peak regions containing the putative PnPf2 binding sites. From this set, overrepresented PWMs up to 20 bp long were modelled with MEME (version 5.1.1) (66, 67). For the resulting PWMs, 500 bp genomic regions centred at ChIP-seq summits were extracted and analysed using CentriMo (Version 5.1.1) to verify that the motif instances were also centred at the respective summits for both the *Pf2-HA* and *Pf2-HA_OE* samples (68). Gene promoters (spanning annotated transcription start sites to the nearest upstream gene feature or 1500 bp) with ≥ 1 occurrence of each motif were determined using FIMO (69). These were cross-referenced with the differentially expressed genes (i.e. expressed significantly up or down in *pf2ko* relative to SN15) defined in a previous RNA-seq analysis (13). Fisher’s exact test with Bonferroni corrected P-values (70) was used to identify *pf2ko* differentially expressed gene-promoter sets significantly enriched (P_adj_ < 0.01) for the respective motifs vs the background rate in SN15.

#### 4.4.3. PnPf2 target gene-promoter analysis

Genes targeted by PnPf2 were determined based on the proximity of summits to annotated genes, which were identified using ChIPseeker (Version 1.24.0) (71). Genes with ≥1 summit falling within their promoter region from the *Pf2-HA* <u>or</u> *Pf2-HA_OE* datasets were considered PnPf2 targets. High-confidence PnPf2 targets corresponded to genes with a promoter summit in *Pf2-HA* <u>and</u> *Pf2-HA_OE*. ChIP-qPCR was then undertaken to verify that the ChIP-seq peak regions in *Pf2-HA* and *Pf2-HA_OE* would also correlate with quantitative enrichment against the *pf2-HA_KO* control strain. Quantitative PCR primer pairs (**File S3**) were designed to flank ChIP-seq summits in a selection of gene promoters (*ToxA*, *Tox1*, *Tox3*, SNOG_03901, SNOG_04486, SNOG_12958, SNOG_15417, SNOG_15429, SNOG_16438, SNOG_20100 and SNOG_30077) and a selection of non-summit control regions (*Act1* and SNOG_15429 coding sequences and the TrpC terminator). The ‘input %’ values were calculated for each sample using the method described previously (72) and used to calculate fold-differences (normalised to *Act1*) for *Pf2-HA* and *Pf2-HA_OE* relative to the *pf2-HA_KO* control for comparison with the *Pf2-HA* and *Pf2-HA_OE* -Log_10_(Q-values) at the respective ChIP-seq summit loci. Pearson’s correlation coefficient was calculated for the ChIP-qPCR fold-difference and ChIP-seq -Log_10_(Q-values) at the respective loci and used as the test statistic to assess whether the association was significant (SPSS version 27.0).

The PnPf2 target genes were cross-referenced with the *pf2ko* expression patterns (expressed significantly up or down in *pf2ko*) defined previously (13) to link direct binding with the modulation of gene expression. The SN15 effector-like genes annotated previously (13) were compiled among the PnPf2 targets. Annotated homologues were identified from the corresponding records in UniProt (release 2020_05) (73). Both the high-confidence and total PnPf2 target-gene sets were then used for GO enrichment/network analysis using the SN15 GO annotations defined previously (13). The ‘enricher’ function in the Clusterprofiler package (Version 3.16.0) (74) was invoked to identify the overrepresented GO classes (P < 0.01) in PnPf2 targets.

### 4.5. Data availability statement

The ChIP-seq reads are available under BioProject ID: PRJNA824526, corresponding to BioSamples SAMN27406642 (*Pf2-HA* strain) and SAMN27406643 (*Pf2-HA_OE* strain).

## 5. Acknowledgements

This study was supported by the Centre for Crop and Disease Management, a joint initiative of Curtin University (https://www.curtin.edu.au/) and the Grains Research and Development Corporation (https://grdc.com.au/) under the research grant CUR00023 Project F3 awarded to KCT). EJ was supported by an Australian Government Research Training Program Scholarship (https://www.dese.gov.au/) administered through Curtin University (https://www.curtin.edu.au/). The funders had no role in study design, data collection and interpretation, or the decision to submit the work for publication.

We would like to acknowledge Dr. Carl Mousley for helpful suggestions relevant to ChIP-seq motif validation, Dr. Darcy Jones for bioinformatics advice and support, as well as Johannes Debbler for assistance with molecular cloning.

