## Supplementary figures and images for "Chromatin-immunoprecipitation reveals the PnPf2 transcriptional network controlling effector-mediated virulence in a fungal pathogen of wheat"

### Fig. S1

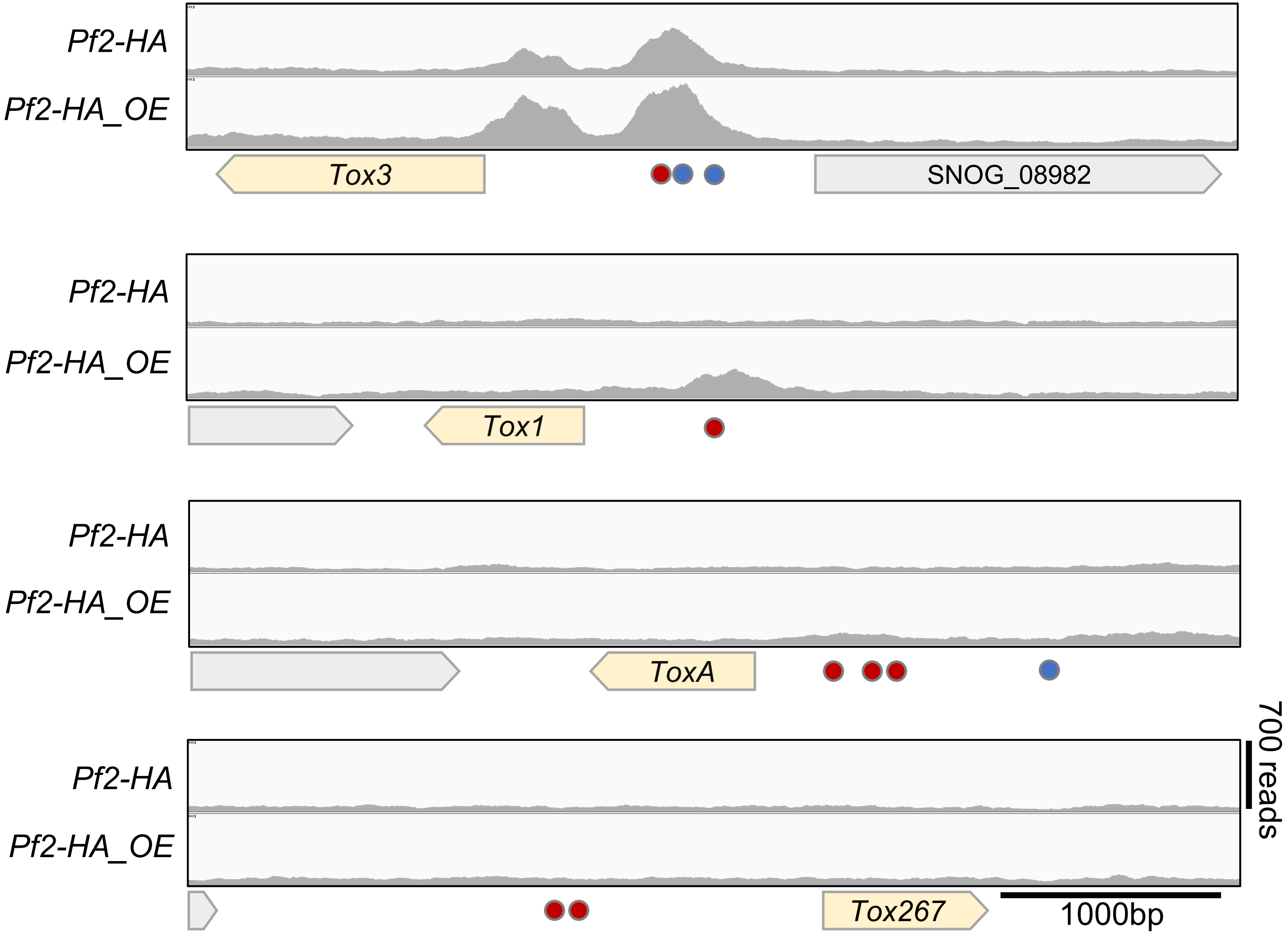

### Fig. S2

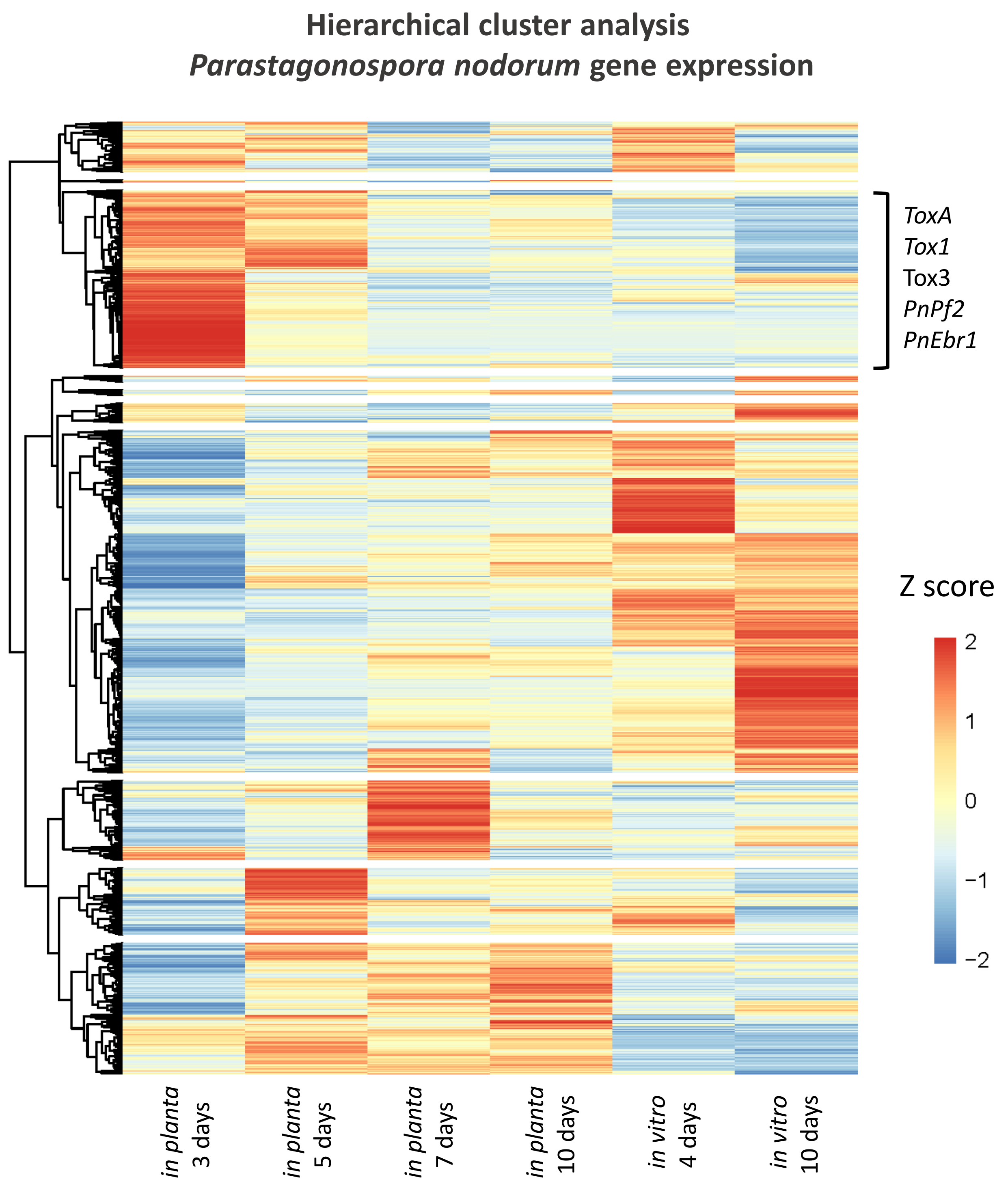
