## Supplementary material for "Chromatin-immunoprecipitation reveals the PnPf2 transcriptional network controlling effector-mediated virulence in a fungal pathogen of wheat": Table S1

**Table S1-A** Effector-like genes predicted as positively regulated direct PnPf2 targets ChIP-seq summit and motif distribution [A]

| Gene ID | | | Motif loci | | Protein length | Protein annotation | Annotated homologues [B] |
|---|---|---|---|---|---|---|---|
| | *Pf2-HA* | *Pf2-HA_OE* | RWMGGVCCGA | CGGCSBBWYYKCGGC | | | |
| *Tox3* | -185;-753 | -189;-760 | -680 | -724; -713; -981 | | | |
| SNOG_13722 | -663 | -668 | - | - | 136 | IPR010829 (Cerato-platanin); IPR009009 (RlpA-like protein, double-psi beta-barrel domain) | *Ds*, *Cb*, *Pf*, *Psf*, *Pt*, *Rc*, *Zb, Zt* |
| SNOG_20100 | -708; -1294 | -697; -1287 | -70; -1303 | -1304; -695 | 71 | - | - |
| SNOG_08150 | - | -204 | -206 | -196 | 124 | - | - |
| SNOG_10510 | -1410 | - | - | - | 346 | - | - |
| SNOG_12449 | - | -179 | - | -984 | 113 | - | *Bm*, *Bo*, *Bs*, *Bv*, *Bz* |
| SNOG_16438 | - | -413 | -507; -1241 | -674 | 138 | - | *Bm*, *Pt* |
| *ToxA* | - | -394 | -216; -366; -409 | -1330 | 178 | IPR021635 (Proteinaceous host-selective toxin ToxA) | *Bs*, *Ptr* |
| *Tox1* | - | -197; -578 | -598 | - | 117 | IPR044057 (Tox1, chitin binding-like domain) | - |
| SNOG_30077 | - | -610 | -609 | - | 67 | - | - |
| SNOG_30352 | - | -189 | - | - | 80 | - | - |

[A] Genes annotated as effectors with significantly reduced expression in the *pf2ko* mutant (1) were considered positively regulated. Identified here as PnPf2 direct-targets based on ChIP-seq summit(s) detected upstream of the start codon in their promoter region, whose relative position is provided along with putative PnPf2 target-motif loci.

[B] Homologous were identified in the respective Uniprot records for: *Bm; Bipolaris maydis, Bo; Bipolaris oryzae, Bs; Bipolaris sorokiniana, Bv Bipolaris victoriae, Bz, Bipolaris zeae, Cb; Cercospora beticola, Pf; Passalora fulva*, *Psf; Pseudocercospora fijiensis, Pt; Pyrenophora teres, Ptr; Pyrenophora tritici-repentis, Rc; Ramularia collo-cygni, Zb; Zymoseptoria brevis, Zt; Zymoseptoria tritici*

**Table S1-B** Transcription factor genes predicted as positively regulated direct PnPf2 targets [A]

| Gene ID | Motif loci | | | | Protein length | Protein annotation | Characterised orthologues [B] |
|---|---|---|---|---|---|---|---|
| | *Pf2-HA* | *Pf2-HA_OE* | RWMGGVCCGA | CGGCSBBWYYKCGGC | | | |
| *PnPro1* (SNOG_03490) | -353; -541 | -270 | - | - | 585 | IPR001138 (Zn2Cys6); IPR021858 (Fun_TF) | AbPro1 (*Ab*), Pro1 (*Cp*), GzZC232 (*Fg*), MoPRO1 (*Mo*), UvPro1 (*Uv*) |
| *PnAda1* (SNOG_04486) | -907 | -903 | -864 | -887 | 654 | IPR004827 (bZIP) | GzbZIP001 *(Fg),* FpAda1 *(Fp),* MobZIP10 *(Mo)* |
| SNOG_08237 | - | -852 | -762 | -853 | 303 | IPR001356 (Homeobox) | CoHox1 (*Co*), MoHox5 (*Mo*), GzHOME004 (*Fg*) |
| SNOG_01243 | -600 | -556 | - | - | 403 | IPR001005 (SANT/Myb) | Myt1 (*Fg*) |
| SNOG_03674 | -382 | -434 | -419 | - | 496 | IPR009071 (HMG box); IPR001660 (SAM) | GzHMG021 (*Fg*) |

[A] Transcription factors with significantly reduced expression in the *pf2ko* mutant (1) were considered positively regulated. Identified here as PnPf2 direct-targets based on ChIP-seq summit(s) detected upstream of the start codon in their promoter region, whose relative position is provided along with putative PnPf2 target-motif loci.

[B] Functionally-characterised orthologues in the scientific literature (2) were identified for: *Ab; Alternaria brassicicola*, *Co; Colletotrichum orbiculare, Cp; Cryphonectria parasitica*, *Fg; Fusarium graminearum, Fp; Fusarium pseudograminearum, Mo; Magnaporthe oryzae*, *Uv; Ustilaginoidea virens*
