## Supplementary material for "Chromatin-immunoprecipitation reveals the PnPf2 transcriptional network controlling effector-mediated virulence in a fungal pathogen of wheat": Table S2

**Table S2** Congruency between genes detected by TF ChIP-seq and RNA-seq in filamentous fungi [A]

| TF studied [B] | ChIP-seq targets | RNA-seq regulated | ChIP-seq + RNA-seq |
|---|---|---|---|
| PnPf2 (*Pn*) **(1)** | 484 | 590 | 93 |
| Clr1 (*Nc*) **(2)** | 164 | 117 | 39 |
| Clr2 (*Nc*) **(2)** | 84 | 132 | 54 |
| Xlr1 (*Nc*) **(2)** | 198 | 90 | 23 |
| Tri6 (*Fg*) **(3)** | 198 | 1614 | 26 |
| FgSR (*Fg*) **(4)** | 119 | 1790 | Not reported |
| Ros1 (*Um*) **(5)** | 1913 | 2006 | 790 |
| MoCrz1 (*Mo*) **(6)** | 346 | 346 (microarray) | 140 |
| CrzA (*Af*) **(7)** | 102 | 3622 | 50 |
| SrbA (*Af*) **(8)** | 97 | 987 | 24 |

[A] Based on the relevant studies in the scientific literature reporting ChIP-seq and RNA-seq differentially-expressed gene datasests.
Abbreviations: TF; transcription factor, ChIP; chromatin immunoprecipitation.
[B] *Af; Aspergillus fumigatus*, *Fg; Fusarium graminearum, Mo; Magnaporthe oryzae*, *Nc; Neurospora crassa, Pn; Parastagonospora nodorum*, *Um; Ustilago maydis*.
