## Supplementary material for "Chromatin-immunoprecipitation reveals the PnPf2 transcriptional network controlling effector-mediated virulence in a fungal pathogen of wheat": Text S1

**Text S1 – Chromatin immunoprecipitation (ChIP) strain assessment and overview of ChIP-seq/ChIP-qPCR.**

This text provides an assessment of the strains used (**Text S1-Fig. 1**), an overview of the ChIP-seq data generation and analysis pipeline (**Text S1-Fig. 2**), the ChIP-seq reads generated for peak/summit calling (**Text S1-Table 1**; **Text S1-Fig. 2**) and the correlation of ChIP-qPCR enrichment with ChIP-seq summit regions representing putative PnPF2 binding loci (**Text S1-Table 2**).

---

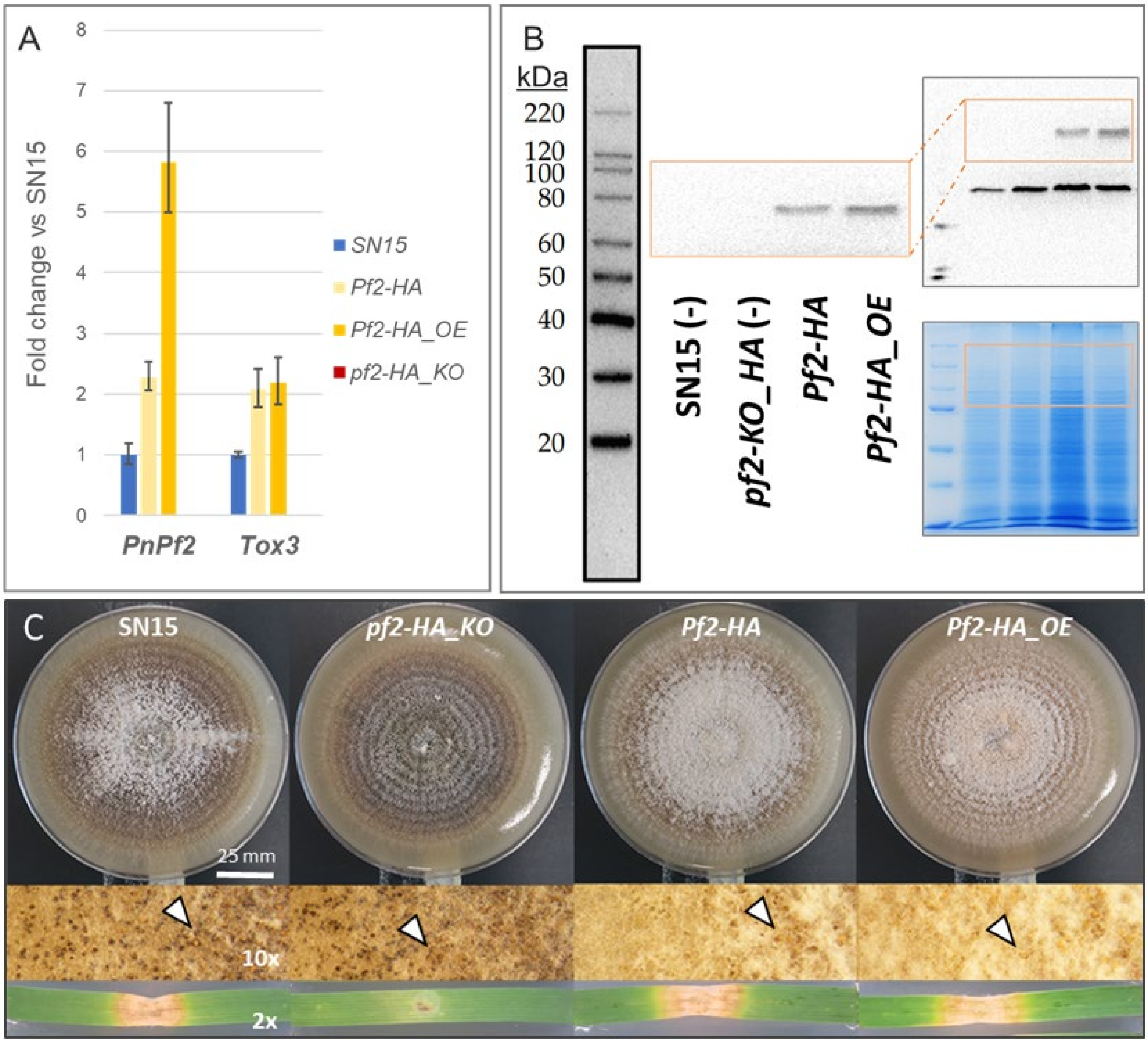


**Text S1-Fig.1** Assessment of strains used for ChIP**.**

**A)** A quantitative PCR analysis that compared the relative gene expression of *PnPf2* and the PnPf2-regulated necrotrophic effector *Tox3* gene. This indicated the 3x haemagglutinin (HA) tag did not inhibit *PnPf2* expression or its activity under the ChIP-seq conditions (Fries3 liquid medium, 72 hrs). Error bars reflect the standard deviation of underlying dCt(*Actin* – *Target*) values. **B)** A Western blot using an anti-HA antibody on total-protein extracts revealed a band corresponding to the expected PnPf2-HA fusion protein size ~75 kDa under ChIP-seq conditions. The band was not detected in SN15 or the negative control strain *pf2-HA_KO*. The full-size original blot and a corresponding Coomassie Blue stained gel are included for reference. A smaller band was detected in all samples presumed to be non-specific antibody binding. **C)** A phenotypic comparison for the respective mutants relative to the SN15 wildtype. The upper two images represent 12 days growth on V8PDA, with arrows indicating mature pycnidia. The lower images depict representative lesions, 12 days following inoculation on detached wheat leaves (cv. Halberd). This suggested virulence was not inhibited and that PnPf2 was functional in *Pf2-HA* and *Pf2-HA_OE* in contrast to *pf2-HA_KO*.

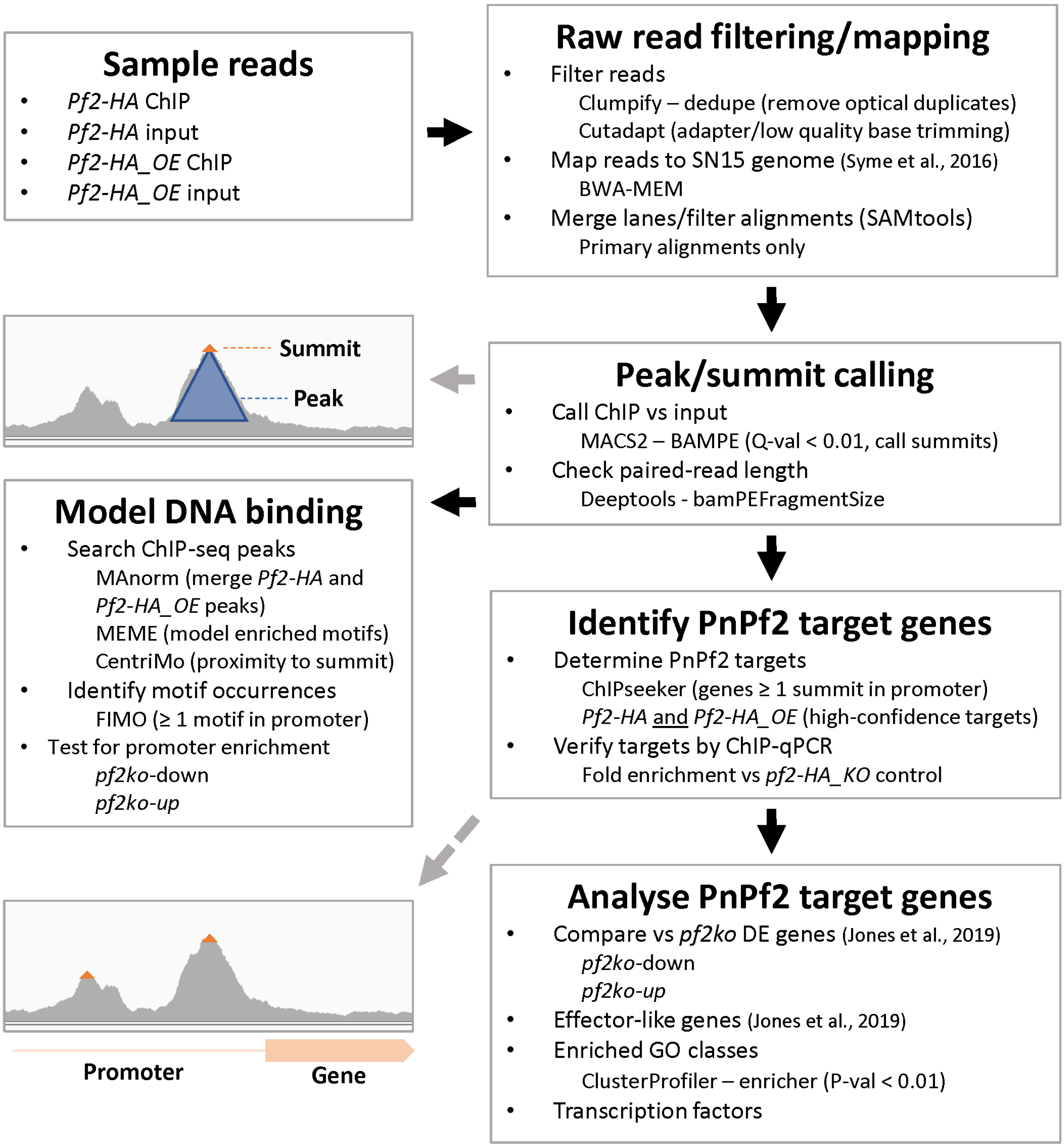


**Text S1-Fig. 2** A diagrammatic overview of the data processing and analysis pipeline followed for ChIP-seq. Grey arrows indicate visual examples of the output from the corresponding stages while black arrows indicate subsequent procedures.

**Text S1-Table 1** ChIP-seq reads mapped to the SN15 genome for calling enriched peaks and summits corresponding to putative PnPf2 binding loci [A]

| Strain | DNA | Raw reads | Mapped pairs | Peak regions | Peak summits |
|---|---|---|---|---|---|
| ***Pf2-HA*** | ChIP | 22,740,293 | 12,416,341 | 740 | 997 |
| | Input | 21,762,558 | 13,307,189 | | |
| ***Pf2-HA_OE*** | ChIP | 29,524,953 | 17,697,729 | 1588 | 2196 |
| | Input | 22,974,742 | 15,270,059 | | |

[A] The raw reads derived from chromatin-immunoprecipitated (ChIP) or background control (Input) for each sample strain. Reads were mapped as pairs to unique loci after quality control and used to call both enriched peak regions and their (one or more) summits.

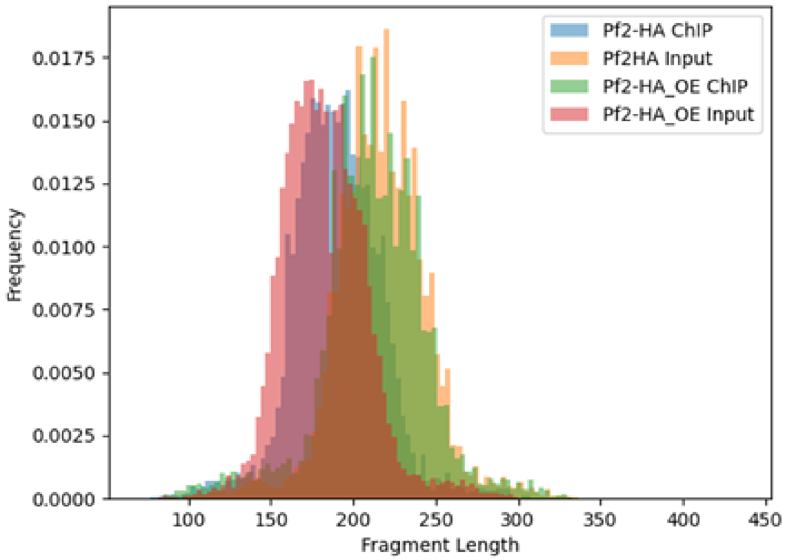


**Text S1-Fig. 3** ChIP-seq paired read length

A summary of the proportional frequency of read-pair lengths that were used for fragment pileup in ChIP and input samples to detect enriched peak/summits from the *Pf2-HA* and *Pf2-HA_OE* strains.

**Text S1-Table 2** ChIP-qPCR relative to ChIP-seq summits at putative PnPf2 binding loci [A]

| Target locus | *Pf2-HA* [B] | | *Pf2-HA_OE* [C] | |
|---|---|---|---|---|
| | qPCR | summit | qPCR | summit |
| *Actin* exon (-) | 1.0 | - | 1.0 | - |
| *TrpC* terminator (-) | 0.9 | - | 1.2 | - |
| *ToxA* promoter | 0.9 | - | 1.4 | 6.5 |
| *Tox3* promoter | 2.2 | 325.9 | 3.7 | 550.7 |
| *Tox1* promoter | 0.8 | - | 3.6 | 121.6 |
| SNOG_03901 promoter | 1.2 | 5.4 | 1.2 | 37.7 |
| SNOG_04486 promoter | 0.9 | 9.5 | 4.4 | 191.2 |
| SNOG_12958 promoter | 2.2 | 236.3 | 5.7 | 252.3 |
| SNOG_15417 promoter | 2.1 | 78.0 | 2.7 | 139.3 |
| SNOG_15429 promoter | 3.5 | 180.4 | 5.9 | 492.9 |
| SNOG_15429 exon (-) | 0.9 | - | 1.7 | - |
| SNOG_16438 promoter | 0.7 | - | 3.4 | 125.4 |
| SNOG_20100 promoter | 1.5 | 21.5 | 3.9 | 160.9 |
| SNOG_30077 promoter | 1.1 | - | 5.4 | 231.0 |

[A] The qPCR values represent fold-enrichment vs the *pf2-HA_KO* control strain and the summit values represent ChIP-seq -$Log_{10}$(Q-values). Target loci listed with (-) were included as qPCR negative controls where no ChIP-seq summit was predicted.

[B] Significantly correlated values (P < 0.01) based on Pearson's correlation (*r* = 0.77).

[C] Significantly correlated values (P < 0.01) based on Pearson's correlation (*r* = 0.74).
