## Supplementary material for "Chromatin-immunoprecipitation reveals the PnPf2 transcriptional network controlling effector-mediated virulence in a fungal pathogen of wheat": Text S2

**Text S2 – Supplemental transcription factor (TF) mutant phenotypic assessment.**

This text provides observations pertaining to the TF mutants generated in this study not presented in the main text. This includes the deletion mutants targeting the putative SNOG_08237 and SNOG_08565 TFs generated in this study (**Text S2-Fig. 1**) and the *PnCreA* mutants relative to *PnPf2* mutants (**Text S2-Fig. 2**).

---

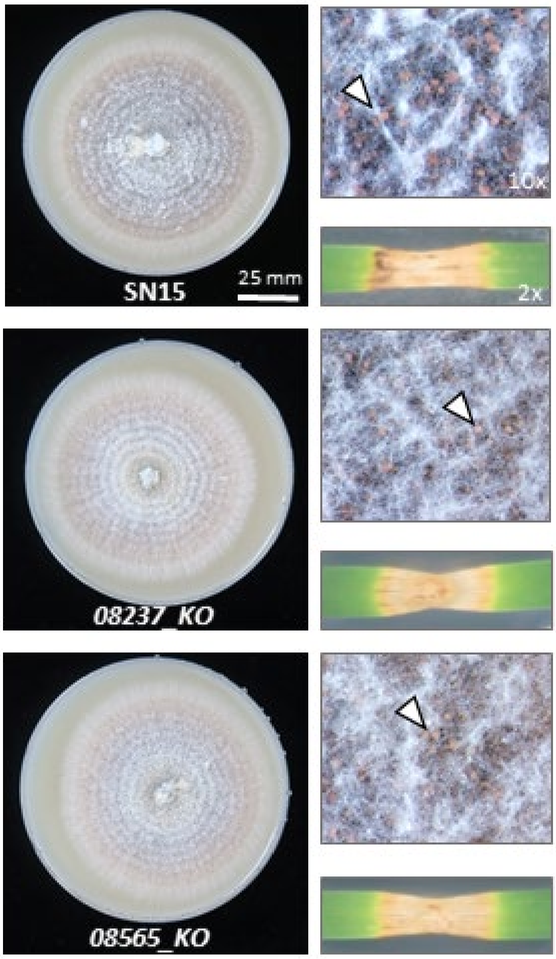


**Text S2-Fig. 1** Assessment of the *08237_KO* and *08565_KO* mutants relative to the SN15 wildtype. Representative images after 12 days of growth on nutrient-rich agar (V8PDA) and infection on detached wheat leaves (cv. Halberd). Arrows demonstrate mature pycnidia detected in the respective mutants.

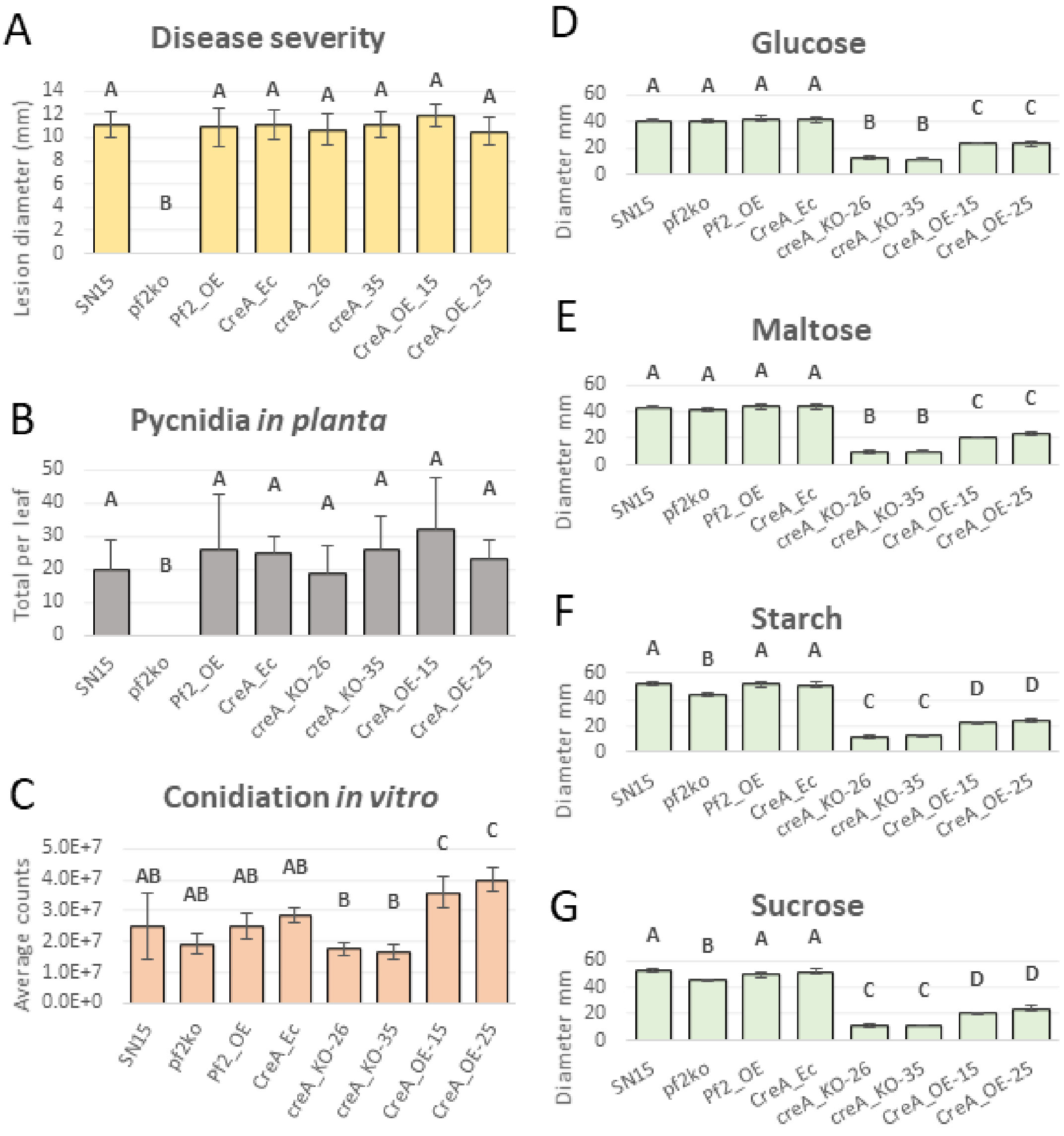


**Text S2-Fig. 2** Phenotypic assessment of *PnCreA* mutants relative to *PnPf2* mutants. **A)** The average lesion sizes representing disease severity and **B)** pycnidia counts, a measure of pathogenic fitness following the infection. **D)** The average conidial (pycnidiospore) counts on V8PDA and **D-G)** the colony diameters following 12 days growth on glucose, maltose, starch or sucrose minimal-medium agar. Letters indicate statistically distinct groupings by ANOVA with Tukey's-HSD ($P<0.05$).
