## Supplementary material for "Chromatin-immunoprecipitation reveals the PnPf2 transcriptional network controlling effector-mediated virulence in a fungal pathogen of wheat": Text S3

**Supplemental Text 3 – Additional materials and methods**

### Contents

DNA cloning ... 2
- Bacterial culturing & transformation ... 2
- Marker templates, PCRs & Sanger sequencing ... 2
- Golden gate and Type IIS cloning ... 2
- Gibson assembly ... 3

General fungal transformation and screening methods ... 5
- Obtaining *PnPf2* and *PnCreA* fungal mutants ... 6
- Obtaining fluorescence-reporter fungal mutants ... 10
- Obtaining additional transcription factor mutants ... 13

Fungal culturing ... 15
- Quantitative PCR for assessing gene expression ... 15
- Protein extraction ... 16
- Western blotting ... 16

Phenotypic analysis of fungal mutants ... 16
- Nuclei staining and microscopy ... 16
- Fungal development ... 17
- Virulence assays ... 17
- Leaf infiltrations using culture filtrate ... 17

Identification of co-expressed transcription factors ... 18

#### DNA cloning

##### Bacterial culturing & transformation

One Shot® TOP10 Chemically Competent *E. coli* cells (Invitrogen, Carlsbad, USA) were used for plasmid transformations following the manufacturers protocol and routinely cultured in SOC media (20 g/L tryptone, 5 g/L yeast extract, 10 mM NaCl, 2.5 mM KCl, 10 mM $MgCl_2$, 10 mM $MgSO_4$, and 20 mM glucose). Plasmid-containing colonies were selected on LB agar (10 g/L tryptone, 5 g/L yeast extract, 10 g/L NaCl, 15 g/L agar) with ampicillin (100 mg/L) or spectinomycin (200 mg/L) and 0.04% v/v X-Gal (Promega, Madison, USA) + 50 µM IPTG (Invitrogen) for blue/white screening when required. Plasmids were extracted using the GenElute Plasmid MiniPrep Kit (Sigma-Aldrich).

##### Marker templates, PCRs & Sanger sequencing

Hygromycin (*HygR*) and Phleomycin (*PhleoR*) resistance-marker templates were derived from the plasmids 'Pan7' and 'Pan8' respectively (1, 2). The *Aspergillus nidulans pGpdA*/*pTef1* promoters and the *tTrpC*/*tTef1* terminator units were derived from 'Pan7' or the 'pFC332' plasmid donated by Uffe Mortensen (3). The *GFP* gene and *pTrpC-HygR-tTrpC* marker cassette were derived from the plasmid 'pGpdGFP' (4). The *dTomato* reporter-gene coding sequence was sourced from a previous publication (5) and the template synthesised (Integrated DNA Technologies, Coralville, Iowa). The 3xHA tag coding sequence was sourced from a synthesised/annealed oligonucleotide pair (**File S3**). For amplification of fragments used in cloning and sequencing, Phusion® High-Fidelity PCR Master Mix (Thermofisher) was used. Fragments were purified using a GenElute™ PCR Clean-Up Kit (Sigma-Adrich, St. Louis, Missouri) or a GenElute™ Gel Extraction Kit where non-specific amplicons were obvious. For size-screening purposes, MyTaq™ DNA Polymerase (Bioline, London, UK) was used for targets up to 5000 bp. Sanger sequencing was performed for sequence-screening purposes on purified plasmids by Macrogen Inc. (Seoul, South Korea) following the recommended guidelines.

##### Golden gate and Type IIS cloning

An in-house Golden Gate (GG) style (6, 7) cloning system was utilised.To summarise, *Bbs*I or *Bsa*I restriction-enzyme sites flank a LacZ marker gene in a pUC19 vector containing either spectinomycin (*SpecR*) or ampicillin (*AmpR*) resistance, termed pGGS-/pGGA-respectively. Four bp overhangs generated by restriction digest with *Bbs*I or *Bsa*I allow *LacZ* marker replacement with the desired fragment sequence(s) in a predetermined orientation, tailored to the production of targeted gene replacement constructs. An overview of the respective fragment overhangs (for the Left flank, Promoter, Coding, Terminator and Right Flank units) used in the directional multi-fragment assembly is provided in **Text S3-Fig. 1**. The one-step digestion/ligation reaction mix consisted 0.08 pmol destination vector, 0.16 pmol fragments (either purified DNA or preassembled in a compatible donor vector), 1.5 µL T4 ligase buffer (Promega), 1.5 µL 10% v/v BSA, 1 µL ATP, 0.5 µL T4 ligase and 0.5 µL either *Bbs*I or *Bsa*I (New England Biolabs, Ipswich, USA) to a total of 20 µL. This was incubated on a 20x cycle (3 min 37 °C then 4 min 16 °C) before enzyme denaturation for 5 min at 80 °C. Where target fragment sequences contained an internal *Bbs*I or *Bsa*I site, 'domestication' was employed (7) to facilitate introduction of synonymous mutations as required. Where required constructs didn't fit the tailored orientation of this Golden Gate system, two-step *Bbs*I (type IIS restriction enzyme) digestion/ligation with custom-designed fragment overhangs was used following the enzyme-manufacturers protocol. Annealed oligonucleotides (produced by mixing 10 µM fragments, heating to 95 °C for 5 min, then cooling at a rate of 1 °C/min until 10 °C) resulting in sequence fragments with appropriate 4 bp overhangs were also incorporated into this system where PCR products were not viable (length too short for PCR or sequence template locally unavailable).

##### Gibson assembly

Where the required fragments were not amenable to TypeIIS restriction enzyme manipulation (such as the presence of multiple internal recognition sequences making 'domestication' unfeasible), Gibson Assembly (New England Biolabs, Ipswich, USA) was used to clone fragments into linear-vector backbones following the manufacturer's protocol.

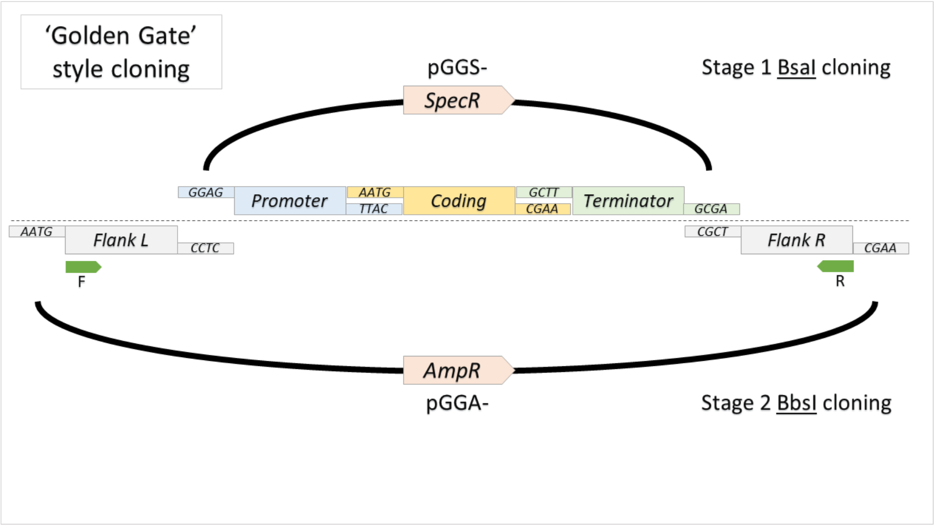

**Text S3-Fig. 1** Overview of Golden Gate style cloning strategy

The universal 4 bp overhangs are indicated for the respective fragments. *SpecR* and *AmpR* refer to the bacterial antibiotic resistance markers for spectinomycin and ampicillin resistance respectively. The first reaction stage incorporating the cloning of fragments into pGGS- is depicted above the dotted line with the second stage assembly incorporating the flanking regions into the destination vector pGGA- underneath. The final assembled construct includes ~700 bp ‘Flanks’ for targeted homologous recombination of the internal modular construct that incorporates gene promoter, coding-sequence (+/- tag) and terminator units. Linear constructs used in subsequent fungal transformations were amplified using primers depicted by the green arrows F/R.

**General fungal transformation and screening methods**

*P. nodorum* transformation was carried out using the polyethylene glycol (PEG) protocol described previously (8). Briefly $10^8$ spores were cultured for 20 hrs in 100 ml CzV8 media (45.4 g/L Czapek Dox liquid media (Oxoid, Bakingstone, UK), 150 ml/L Campbell's V8 juice, 20.0 g/L casamino acids, 20 g/L peptone, 20 g/L yeast extract, 3 g/L adenine, 0.02 g/L biotin, 0.02 g/L nicotinic acid, 0.02 g/L p-aminobenzoic acid, 0.02 g/L pyridoxine and 0.02 g/L thiamine) at 100 rpm and 22 $^{o}$C in the dark on an orbital shaker. Mycelia were treated with 15mg/mL Extralyse (Laffort, Floirac, France) + 1.2 M $MgSO_4$ for 2 hrs at 28 $^{o}$C to form protoplasts. Five µg of linear DNA construct suspended in STC buffer (1 M sorbitol, 10 mM Tris-HCl, 10 mM $CaCl_2$) was added to 100 µL protoplasts (washed in 2 M sorbitol, resuspended in 1 mL STC buffer) and incubated for 20 min in 60% PEG. Successful transformants were selected for in Hygromycin (200mg/L) or Phleomycin (50 mg/L) supplemented CzV8 agar (CzV8 + 10 g/L agar and 18 g/L sorbitol) before subculturing onto individual V8PDA plates containing the respective antibiotics to obtain mutant-fungal material for both DNA extraction with a Biosprint DNA Plant Kit (QIAGEN, Hilden, Germany) and long-term storage in 20% glycerol at -80 $^{o}$C.

Mutants were selected for subsequent analysis through a combination of PCR and qPCR gDNA-screening approaches. The PCR screening utilised primers (**File S3**) that flank the respective targeted integration sites to ensure that constructs had incorporated at the desired locus. A robust qPCR-based method was used to verify a single construct had been integrated into the genome, which followed a method previously described (9). Briefly, a standard curve of Starting Quantity (SQ) values was generated from serially diluted reference gDNA (5 ng/µL to 0.16 ng/µL). Reference gDNA was sourced from SN15 or the *pf2ko* mutant (10, 11) and the Pf2_qPCR_F/R or the Phleo_qPCR_F/R primer-pairs utilised respectively. The SN15 reference was applied for gene-replacement candidate mutants while the *pf2ko* reference was applied for gene-deletion candidates (described in subsequent paragraphs). The Actin_qPCR_F/R primer-pair targeted the *Act1* gene as an internal-normalisation standard. Sample reactions were performed in triplicates at 1.6 ng/µL gDNA and an average SQ ratio (*Act1*/*PnPf2* or *Act1*/*PhleoR*) between 0.8 and 1.2 was considered a single copy.

**Obtaining *PnPf2* and *PnCreA* fungal mutants**

A *PnPf2* knockout (KO) construct was first produced through GG cloning by attaching ~700 bp universal *PnPf2* flanking-fragments located 5' and 3' of *PnPf2* (amplified from SN15 gDNA with Pf2_HR_FL_BsaI_F/R and Pf2_HR_FR_BsaI_F/R respectively) to the *pTef1-PhleoR-tTef1* marker (**Text S3-Fig. 2A**). The resulting construct was amplified with Pf2_HR_FL_F/Pf2_HR_FR_R which was then used to generate the new *pnpf2* mutant (*pf2_KO*) from the wildtype (SN15) by PEG transformation. The *pf2_KO* mutant was amenable to homologous recombination (HR) and marker retrieval at the native locus, which is where *PnPf2* tagged constructs were subsequently introduced (using the *pTrpC-HygR-tTrpC* selectable marker).

The *PnPf2-GFP* construct was produced by first amplifying the *PnPf2* region with the pPf2_P_BbsI_985_F/Pf2_link_B_BbsI_R primer-pair from SN15 (encompassing the promoter, coding sequence and incorporating a GGSG peptide linker for protein/tag spatial separation) followed by Type IIS cloning into the linear vector amplified from 'pGpdGFP' (4) using eGFP_BbsI_F/pBack_FL_BbsI_R. The resulting construct was PCR amplified using tTrpc_T_BbsI_F/eGFP_B_BbsI_R to receive the *PnPf2* terminator sequence amplified from SN15 using tPf2_T_BbsI_F/R by Type IIS cloning to yield the plasmid 'Pf2-GFP_HygR' (**Text S3-Fig. 2B**). Two separate linear vectors were amplified from 'Pf2-GFP_HygR' using either Pf2_link_B_BbsI_R/tPf2_T_BbsI_F or pPf2_P_BbsI_R/tPf2_T_BbsI_F. These vectors were used for incorporating the oligo-annealed HA_Oligo_sense/anti fragment (encoding a 3x haemagglutinin tag) by TypeIIS cloning to produce the plasmids 'Pf2-HA_HygR' and 'pf2-HA_KO_HygR' respectively (**Text S3-Fig. 2C**). Linear constructs were amplified from 'Pf2-GFP_HygR', 'Pf2-HA_HygR and 'pf2-HA_KO_HygR' using pPf2_P_BbsI_985_F/pTrpC_T_BbsI_R to attach the same ~700 bp 5' and 3' *PnPf2* flanking fragments (used for *pf2_KO*) by GG cloning. The resulting plasmids were then amplified with Pf2_HR_FL_F/Pf2_HR_FR_R to obtain the linear constructs used for HR by PEG transformation in the *pf2_KO* background to generate the fungal mutants *Pf2-GFP*, *Pf2-HA* and *pf2-HA_KO* respectively (**Text S3-Fig. 2E**).

*PnPf2* overexpression mutants were derived using a promoter replacement strategy where a *pGpdA* promoter was used to replace the native *PnPf2* promoter. The replacement construct was produced through GG cloning, where the same ~700 bp 5' *PnPf2* left-flanking fragment (used for *pf2_KO*) was used as well as a 3' right-flanking fragment derived from two

PCR products. The two products, designed to assemble an in-frame overexpression unit, were amplified using pGpd_FR_BsaI_F/R from the 'Pan7' plasmid (2) and Pf2_OE_FR_BsaI_F/R from SN15 gDNA. The respective flanks were fused either side of the recycled *pTef1-PhleoR-tTef1* marker in the GG reaction (**Text S3-Fig. 2D**). The resulting construct was amplified using Pf2_HR_FL_F/Pf2_mid_FR_R to produce the linear constructs for HR by PEG transformation in the SN15, *Pf2-GFP* and *Pf2-HA* backgrounds, producing the strains *Pf2_OE*, *Pf2-GFP_OE* and *Pf2-HA_OE* respectively (**Text S3-Fig. 2E**).

The *PnCreA* KO and overexpression mutants were derived in an analogous procedure to the respective *PnPf2* mutants *pf2_KO* (**Text S3-Fig. 2A**) and *Pf2_OE* (**Text S3-Fig. 2D**). A common *PnCreA* 5' left-flanking fragment was amplified from SN15 gDNA using the primer-pair CreA_HR_FL_BsaI_F/R. The 3' right-flanking unit for KO was amplified using CreA_HR_FR_BsaI_F/R. The 3' right-flanking overexpression unit was formed from the fragment amplified with pGpd_FR_BsaI_F/R from 'Pan7' and the fragment amplified with CreA_OE_FR_BsaI_F/R from SN15 gDNA. The common left-flanking unit was fused to the *pTef1-PhleoR-tTef1* marker in the GG reactions that incorporated the right-flanking KO or overexpression units to generate the KO and overexpression plasmid-constructs respectively. The linear KO construct was amplified using the primer-pair CreA_HR_FL_F/R and the overexpression construct amplified using the primer-pair CreA_HR_FL_F/ CreA_OE_FR_R, both of which were separately integrated by HR into the SN15 background by PEG transformation to produce the mutants *creA_KO* and *CreA_OE* respectively (**Text S3-Fig. 2E**). An ectopically integrated KO-construct mutant (*CreA_Ec*) was also retained as a control for subsequent phenotypic analyses.

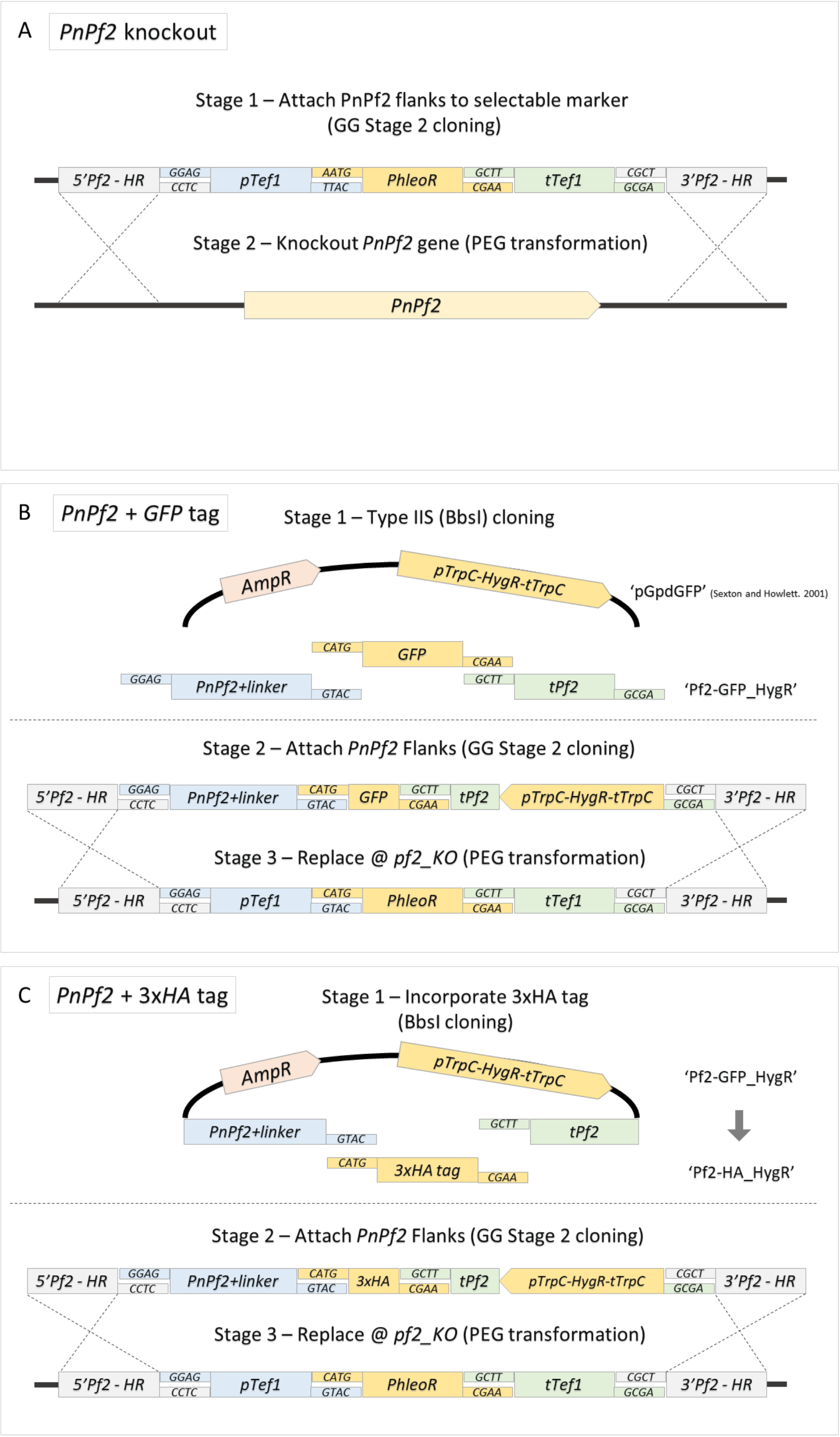
A
PnPf2 knockout
Stage 1 – Attach PnPf2 flanks to selectable marker
(GG Stage 2 cloning)
5'Pf2 - HR
GGAG
CCTC
pTef1
AATG
TTAC
PhleoR
GCTT
CGAA
tTef1
CGCT
GCGA
3'Pf2 - HR
Stage 2 – Knockout PnPf2 gene (PEG transformation)
PnPf2
B
PnPf2 + GFP tag
Stage 1 – Type IIS (BbsI) cloning
AmpR
pTrpC-HygR-tTrpC
'pGpdGFP' (Sexton and Howlett. 2001)
CATG
GFP
CGAA
GGAG
PnPf2+linker
GTAC
GCTT
tPf2
GCGA
'Pf2-GFP_HygR'
Stage 2 – Attach PnPf2 Flanks (GG Stage 2 cloning)
5'Pf2 - HR
GGAG
CCTC
PnPf2+linker
CATG
GTAC
GFP
GCTT
CGAA
tPf2
pTrpC-HygR-tTrpC
CGCT
GCGA
3'Pf2 - HR
Stage 3 – Replace @ pf2_KO (PEG transformation)
5'Pf2 - HR
GGAG
CCTC
pTef1
CATG
GTAC
PhleoR
GCTT
CGAA
tTef1
CGCT
GCGA
3'Pf2 - HR
C
PnPf2 + 3xHA tag
Stage 1 – Incorporate 3xHA tag
(BbsI cloning)
AmpR
pTrpC-HygR-tTrpC
'Pf2-GFP_HygR'
PnPf2+linker
GTAC
GCTT
tPf2
CATG
3xHA tag
CGAA
'Pf2-HA_HygR'
Stage 2 – Attach PnPf2 Flanks (GG Stage 2 cloning)
5'Pf2 - HR
GGAG
CCTC
PnPf2+linker
CATG
GTAC
3xHA
GCTT
CGAA
tPf2
pTrpC-HygR-tTrpC
CGCT
GCGA
3'Pf2 - HR
Stage 3 – Replace @ pf2_KO (PEG transformation)
5'Pf2 - HR
GGAG
CCTC
pTef1
CATG
GTAC
PhleoR
GCTT
CGAA
tTef1
CGCT
GCGA
3'Pf2 - HR

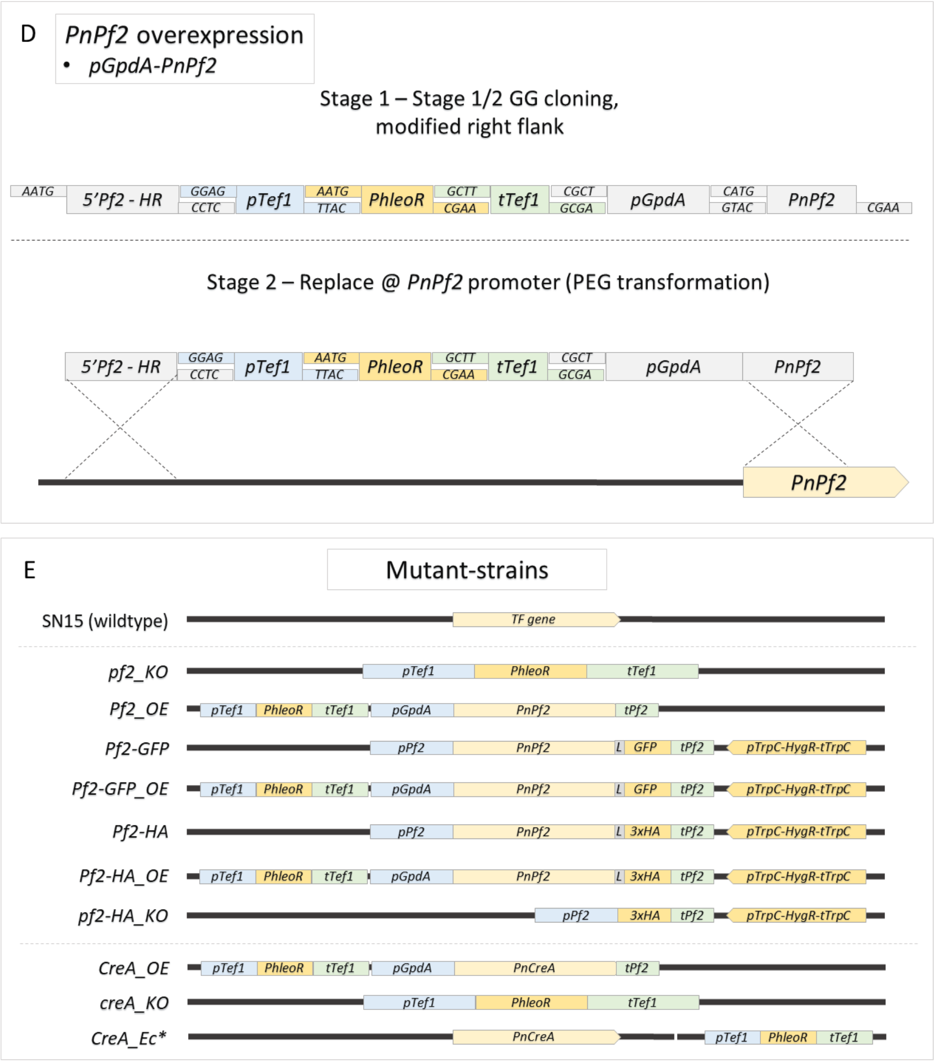


**Text S3-Fig. 2** *PnPf2* and *PnCreA* mutants

An overview of the cloning and transformation stages used to produce the *PnPf2* and *PnCreA* mutants assessed in this study. Panel **A** depicts the generation of the *pf2_KO* mutant that formed the basis for reintroduction (and *PhleoR*-marker retrieval) of the *GFP* and 3x*HA* tagged *PnPf2* constructs to produce the *Pf2-GFP* and *Pf2-HA* mutants depicted in panel **B** and **C** respectively. The *Pf2_OE*, *Pf2-GFP_OE* and *Pf2-HA_OE* strains were produced by introducing the overexpression construct depicted in panel **D** into the wildtype SN15, *Pf2-GFP* and *Pf2-HA* strains respectively. Overexpression was driven by the *pGpdA* promoter. The *CreA_OE* and *creA_KO* mutants were generated analogous to the respective *Pf2_OE* and *pf2_KO* mutants, but targeted to the *PnCreA* gene locus in the wildtype background. Panel **E** provides and overview of the strains produced relative to the wildtype gene loci for *PnPf2* and *PnCreA* mutants. *Indicates non-targeted (ectopic) integration of the respective constructs.

**Obtaining fluorescence-reporter fungal mutants**

A strain for epifluorescence microscopy was produced in the SN15 background. To this end, a *pTef1-GFP-tTef1* expression cassette was first derived by GG cloning which produced the plasmid 'pGGS_pTef1-GFP' (**Text S3-Fig. 3A**). From this a linear plasmid was amplified using the primer-pair pTef1_F/pGG_Gibson_R, which was used to incorporate the *pGpdA-PhleoR-tTrpC* cassette (amplified from 'Pan8' with PhleoR_Gibson_F/R) by Gibson assembly. A linear construct was amplified from the resulting plasmid 'pTef1-GFP_PhleoR' using pGG_screen_F/R and integrated into SN15 by PEG transformation to obtain the mutant SN15-*GFP* (**Text S3-Fig. 3D**).

A strain constitutively expressing the *dTomato* reporter gene was produced in the SN15 background. To this end, the *pGpdA-PhleoR-tTrpC* cassette was first incorporated upstream of the *pTef1-dTomato-tTef1* reporter-gene cassette into the 'pGGS-' plasmid backbone by GG cloning. For the cloning reaction, the promoter fragment was amplified' using the primer-pair pGpd_P_BbsI_F/pTef1_P_BbsI_R from the 'pTef1-GFP_PhleoR' plasmid template, which positioned the *pGpdA-PhleoR-tTrpC* directly upstream of the *pTef1* promoter in the *pTef1-dTomato-tTef1* cassette (**Text S3-Fig. 3B**). A 5' left-flank and a 3' right-flank were then attached to the *pGpdA-PhleoR-tTrpC-pTef1-dTomato-tTef1* double-cassette in the 'pGGA-' plasmid backbone by GG cloning to produce the plasmid 'pUbc6_pTef1-dTom' (**Text S3-Fig. 3B**). The respective 5' and 3' flanks were amplified from SN15 gDNA using the primer-pairs Ubc6_HR_FL_BsaI_F/R and Ubc6_HR_FR_BsaI_F/R to facilitate targeted integration at a predefined locus. A linear construct was amplified from 'pUbc6_pTef1-dTom' using the primer-pair Ubc6_HR_FL_F/R and incorporated via PEG transformation into the SN15 background as a single copy to obtain the mutant *pTef1-dTom* (**Text S3-Fig. 3D**). The integration locus was downstream of *PnUbc6* (12), a conserved homologue of the *Saccharomyces cerevisiae* housekeeping/stably-expressed reference gene *UBC6* (13).

The 'pUbc6_pTef1-dTom' plasmid was used as a template to amplify a linear-backbone with the primer-pair dTom_Gibson_F/tTrpC_Gibson_R. The resulting backbone 'pUbc6-dTomato' could receive the *SNOG_15417* promoter (*p15417*) in place of *pTef1* by Gibson assembly (**Text S3-Fig. 3C**). Two novel *p15417* mutations were first introduced either alone, or in combination by the GG cloning 'domestication' procedure (7) in a 'pGGA-' plasmid backbone. The first mutation (*m1*) was assembled using two amplicons derived from SN15 gDNA using the primer-pairs p15417_BsaI_F/p15417_M1_BsaI_R and p15417_m1_BsaI_F/p15417_BsaI_R. The second mutation (*m2*) was assembled using two PCR products amplified using p15417_BsaI_F/p15417_m2_BsaI_R and p15417_m2_BsaI_F/p15417_BsaI_R. A combination of the mutations (*m1m2*) was introduced by assembling three amplicons derived with the primer-pairs p15417_BsaI_F/p15417_m2_BsaI_R, p15417_m2_BsaI_F/p15417_m1_BsaI_R and p15417_m1_BsaI_F/p15417_BsaI_R (**Text S3-Fig. 3C**). The resultant mutated promoters, as well as a non-mutated promoter (*M1M2*), were amplified using the primer-pair p15417_Gibson_F/R, which formed the respective linear fragments used for Gibson assembly into the 'pUbc6-dTomato' linear backbone in-frame of the *dTomato* coding sequence. From the resultant constructs, the primer-pair Ubc6_HR_FL_F/R was used to amplify and integrate the respective linear constructs by HR through PEG transformation in the SN15 background to obtain the mutants *p15417_M1M2*, *p15417_m1M2*, *p15417_M1m2* and *p15417_m1m2* (**Text S3-Fig. 3D**).

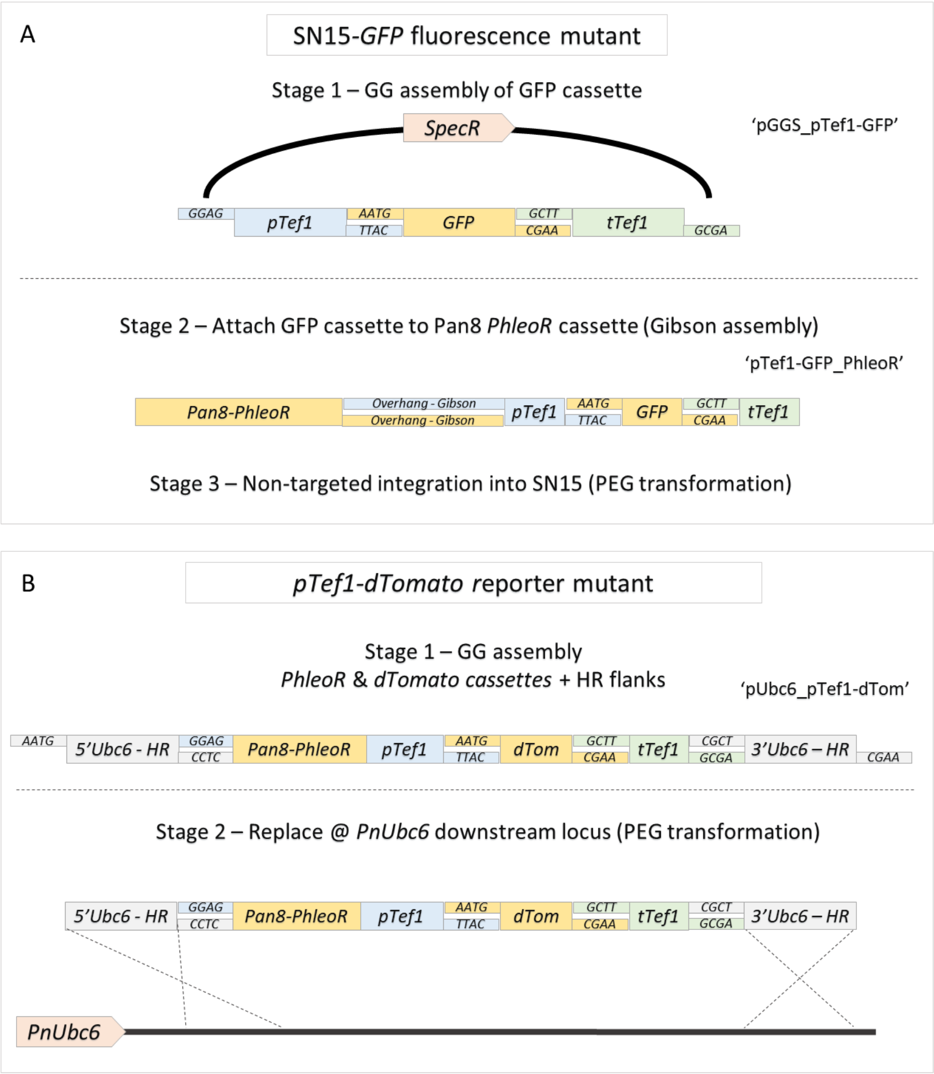

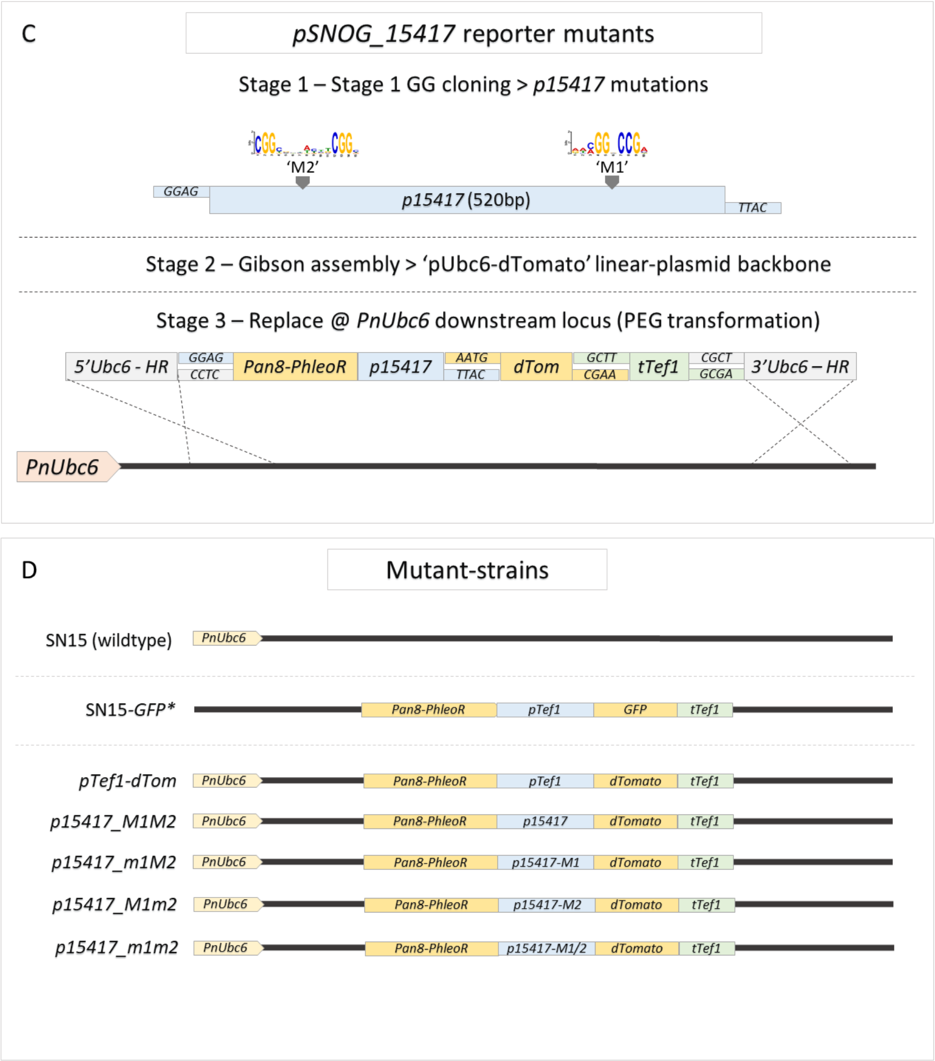


**Text S3-Fig. 3** Fluorescence reporter mutants

An overview of the cloning and transformation stages used to produce the fluorescence reporter mutants used in this study. Panel **A** depicts the generation of the SN15-*GFP* mutant used for microscopic analysis where *GFP* was driven by the *Tef1* promoter. Panel **B** depicts the generation of the *pTef1-dTomato* mutant, where the *dTomato* reporter gene driven by the *Tef1* promoter was integrated as a single copy at the predefined locus downstream of *PnUbc6*. This locus was also used to integrate the SNOG_15417 promoter-*dTomato* reporter constructs outlined in panel **C**, where promoter mutations at Motif 1 (*m1*) and/or Motif 2 (*m2*) were introduced. Panel **D** provides an overview of the strains produced relative to the wildtype gene loci for *PnPf2* and *PnCreA* mutants. *Indicates non-targeted (ectopic) integration of the respective constructs.

**Obtaining additional transcription factor mutants**

The SNOG_03067 (*PnEbr1*), SNOG_03490 (*PnPro1*), SNOG_04486 (*PnAda1*), SNOG_08237 and SNOG_08565 TF KO mutants were generated in the SN15 background. The HR gene KO constructs were first derived by attaching 5' and 3' flanks (amplified from SN15 using the primer-pairs *TF*_HR_FL_BsaI_F/R and *TF*_HR_FR_BsaI_F/R; *TF* corresponds to the numerical SNOG_ID for the respective gene annotations) to the *pGpdA-PhleoR-tTrpC* cassette by GG cloning (**Text S3-Fig. 4A**). The linear constructs were amplified from the resulting plasmids using *TF*_HR_FL_F/*TF*_HR_FR_R and used for PEG transformation to obtain the KO mutants *ebr1_KO*, *pro1_KO*, *ada1_KO*, *08237_KO* and *08565_KO* (**Text S3-Fig. 4B**). For gene complementation, the original TF genes were amplified from SN15 gDNA using *TF*_Gibson_F/R. A plasmid 'pGGS_pTef1-HygR' was then assembled via GG cloning that contained the *pTef1-HygR-tTef1* marker. This was amplified using the primer-pair pTef1_F/pGG_Gibson_R, to form a linear backbone an integrate the respective TF-gene amplicons by Gibson assembly (**Text S3-Fig. 4A**). The resulting constructs were amplified from the respective plasmids using pGG_screen_F/R and randomly integrated into one of the respective KO mutant backgrounds by PEG transformation to produce the complemented mutants (**Text S3-Fig. 4B**).

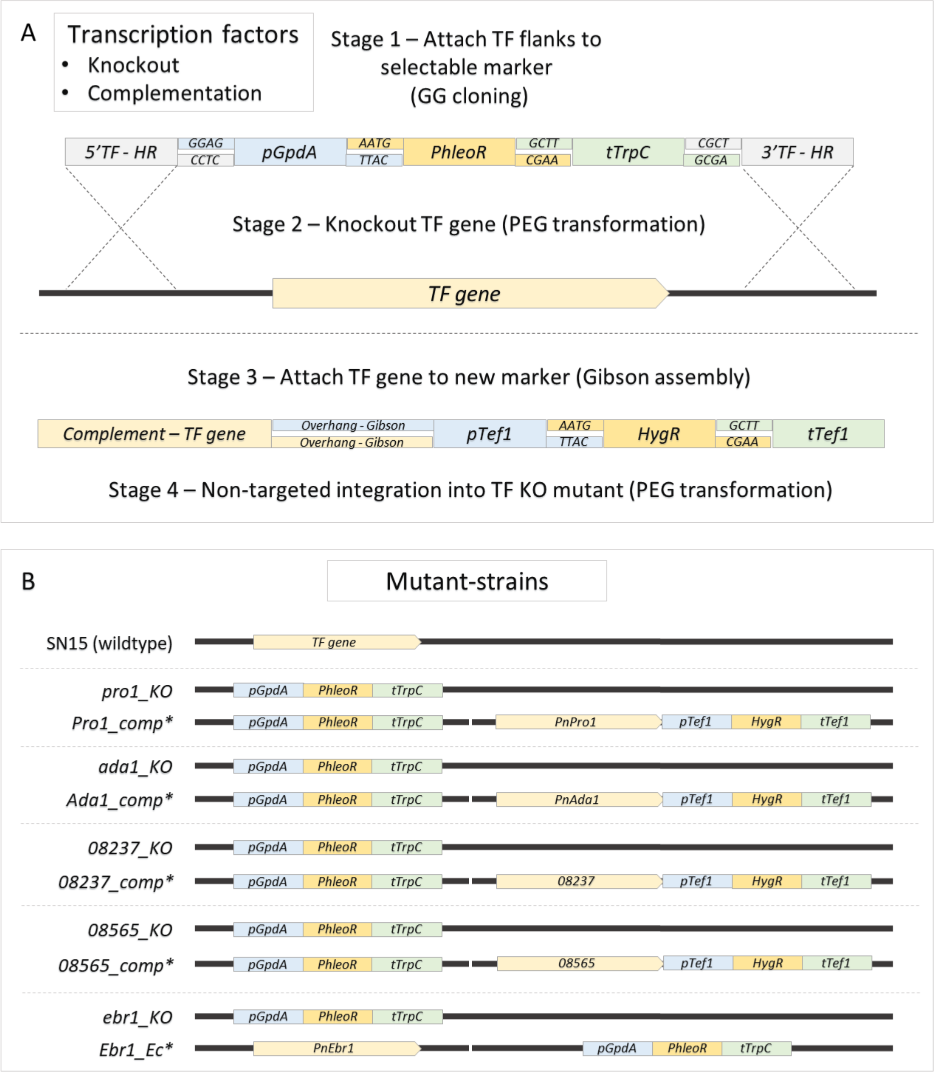


**Text S3-Fig. 4** Additional transcription factor mutants

An overview of the cloning and transformation stages used to produce the additional TF mutants used in this study. Panel **A** provides a general depictions of the analogous strategy used to generate the respective constructs for gene deletion through HR, and then to complement the respective TF genes into the KO strain. Panel **B** depicts provides an overview of the strains produced relative to the wildtype gene loci the respective mutants. *Indicates non-targeted (ectopic) integration of the respective constructs.

**Fungal culturing**

Stock cultures of the *P. nodorum* wildtype SN15, and the derived mutants, were kept in 20% glycerol at -80 °C and periodically inoculated to sterile V8PDA plates (10 g/L potato dextrose agar, 3 g/L CaCl2, 150 mL/L V8 juice and 15 g/L w/v agar) for 12 days under a 12 hr fluorescent light/dark cycle for general use. Pycnidiospores were collected for inoculation in subsequent assays as previously described (14). Sterilised Fries3 medium was used for general liquid culturing following previous studies (15, 11) which consisted 5 g/L ammonium tartrate ((NH4)2C4H4O6), 1 g/L NH4NO3, 50 mg/L MgSO4.7H2O, 130 mg/L $K_2HPO_4$, 260 mg/L $KH_2PO_4$, 30 g/L sucrose, 1 g/L yeast extract and 2 mL/L trace stock (167 mg/L LiCl, 227 mg/L $CuSO_4.5H_2O$, 34 mg/L $H_2MoO_4$, 72 mg/L $MnCl_2.4H_2O$ and 80 mg/L $CoCl_2.4H_2O$ in $H_2O$) in $H_2O$. Mycelia were grown in Fries3 by inoculating spores to a standardised final concentration of $10^4$ $mL^{-1}$ and grown in the dark for 72 hrs, 22 °C and 100 rpm on an orbital shaker prior to downstream analyses.

**Quantitative PCR for assessing gene expression**

Mycelia were grown in 2 mL Fries3 (see ‘Fungal culturing’ above) in 12 well microtiter plates (Corning, Somerville, USA) before transfer into 2 mL safelock tubes (Sarstedt, Nümbrecht, Germany) for collection via centrifugation (12000 g, 1 min). Pellets were then washed once with Milli-Q® purified $H_2O$ before snap-freezing/lyophilisation for 12 hrs in a freeze dryer (Zirbus, Bad Grund, Germany). Mycelia was then crushed using tungsten beads with a TissueLyser II (QIAGEN, Hilden, Germany) before RNA extraction using a PureLink RNA Mini Kit (Thermofisher, Waltham, USA), DNAse treatment (DNA-free DNA Removal Kit, Thermofisher) and cDNA synthesis (iScript cDNA Synthesis Kit, Biorad, Hercules, USA). Quantitative PCR was undertaken on a Biorad CFX96™ thermocycler using the RT-PCR SYBR® Green master mix (QIAGEN, Hilden, Germany). Reactions were performed using 600 nM primers and 2 ng/µL cDNA in a 20 µL solution. 38 cycles were undertaken (95 °C 15 sec, 58 °C 30 sec, 72 °C 30 sec) followed by a melt curve analysis step using the Precision Melt Analysis™ software (Biorad) following the manufacturer's guidelines to ensure primer specificity. The *PnPf2*, *Tox3*, *ToxA*, *PnCreA* and *dTomato* genes were targeted using the Pf2_qPCR_F/R, Tox3_qPCR_F/R, ToxA_qPCR_F/R, CreA_qPCR_F/R or dTomato_qPCR_F/R primer-pairs respectively. The relative abundance of target-gene cDNA was calculated with the $2^{dCt}$ method (16) using *Act1* amplified with Actin_qPCR_F/R primers as the internal standard, which is regularly used in *P. nodorum* (17, 10, 18). Fold differences between samples were calculated as $2^{ddCt}$ values.

##### Protein extraction

Mycelia were grown in 100 mL Fries3 (see ‘Fungal culturing’ above) in a sterile 250mL conical flask. For protein extractions, the mycelia were instead transferred to 50 mL Falcon tubes for collection via centrifugation (5 min, 3000 g) before washing 1x with Milli-Q® purified $H_2O$ and snap-freezing/lyophilisation in a freeze-dryer (Zirbus, Harz, Germany) for at least 24 hrs. Mycelia were then weighed and crushed in liquid nitrogen using a mortar/pestle before the addition of 10x (v/w) ice-cold lysis buffer (50 mM Tris, 150 mM NaCl, 1 mM EDTA and 10 mL/L TritonX100 with pH = 8. Additionally, 0.1% v/v NaDOC, 1 mM PMSF and 1% v/v protease inhibitor cocktail (P8215, Sigma-Aldrich, St Louis, USA) were freshly added before use). 1.6 mL of resuspended material was then transferred to a sterile 2 mL tube before gentle rotation at 4 °C for 20 min. Protein fractions were then separated by taking 1mL supernatant from two rounds of centrifugation (5000 g, 4 °C for 5 min).

##### Western blotting

Protein preparations were quantified using a Direct Detect infrared spectrometer (Merck, Kenilworth, USA) prior to downstream analysis. Western blotting was undertaken on whole protein extracts, probed using an anti-HA polyclonal antibody (71-5500 - Thermofisher, Waltham, Massachusetts) and detected using an anti-IgG/Pierce ECL chemiluminescence detection system (A16096/32209 - Thermofisher).

#### Phenotypic analysis of fungal mutants

##### Nuclei staining and microscopy

Pycnidiospores of SN15 and the fluorescence strains SN15-*GFP* and *Pf2-GFP_OE* were inoculated onto 1% potato dextrose agar set on microscope slides and left for 24 hrs in the dark in a humid container to germinate. Fungal material was flooded with a fixing solution (4% w/v formaldehyde and 0.1% Triton in 1x PBS; phosphate buffered saline at pH 7.4) for 10 min, then continuously washed with a rinsing solution (0.1% Triton/PBS) for 10 sec, stained for 10 min (0.1% Triton/PBS with 0.5µg/mL DAPI; 4′,6-diamidino-2-phenylindole) and washed again with the rinsing solution followed by a 1x PBS wash before a coverslip was applied. Samples were examined on an Olympus BX-51 microscope with a 40x objective lens and a DAPI (Ex 350/50, FT 400, BP 460/50) or FITC (Ex 480/30, FT 505, BP 535/40) filter for visualisation of stained nuclei and GFP fluorescence, respectively.

##### Fungal development

Fungal development was assessed following 12 days growth, after inoculating 10 µL of $10^6$ spores $mL^{-1}$ on various media. The conidiation rate was assessed on V8PDA *in vitro* by counting pycnidiospores in three replicates. Oxidative stress inhibition was measured by comparing the radial growth of strains on minimal-medium (MM) agar plates (10 g sucrose, 2 g $NaNO_3$, 1 g $K_2HPO_4$, 0.5 g KCl, 0.5 g $MgSO_47H_2O$, 0.01 g $ZnSO_47H_2O$, 0.01 g $FeSO_47H_2O$ and 2.5 mg $CuSO_45H_2O$ $L^{-1}$) with and without 20 mM $H_2O_2$. A relative measure of fitness was obtained by dividing the diameter with/without $H_20_2$ across three replicates. For both assays, a one-way ANOVA with Tukeys-HSD post-hoc test was used to test for differences ($P < 0.05$) between strains (SPSS version 27.0). Defects in carbon-source utilisation were assessed through growth on MM agar, where sucrose was substituted with 10 g/L glucose, maltose and starch. The starch-MM agar plates were post-stained with Lugol's iodine to identify zones of hydrolysis.

##### Virulence assays

Wheat seedlings were grown for 12 days in vermiculite supplemented with minimal amounts of all-purpose fertiliser (Yates, Auckland, New Zealand) under a 12 hr light/dark photoperiod at 22 °C in a controlled growth chamber (Conviron, Winnipeg, Canada) before subsequent use. For the assessment of fungal virulence, the detached leaf assay (DLA) was used (19). Five cm excisions from the first leaves of wheat seedlings were embedded in 75 mg/L benzimidazole agar containing plates. A 10 µL inoculation of $10^6$ spores $mL^{-1}$ in 0.02% Tween was applied to the centre of each leaf (if non-sporulating mutants were assessed, 3 mm-diameter mycelial plugs were used) before returning to the growth chamber. Virulence was quantified using DLAs after 12 days with 10 replicates (5x Halberd and 5x Calingiri) using two metrics; lesion diameter (mm) and weighted pycnidia counts (scored as immature/black = 1, mature/pink = 2, fully mature/burst = 3). For both measures, a one-way ANOVA with Tukeys-HSD post-hoc test was used to test for differences ($P < 0.05$).

##### Leaf infiltrations using culture filtrate

First leaves on 12-day old wheat seedlings were infiltrated with culture filtrate following a method described previously (20). For the production of culture filtrate, 100mL cultures of mycelia grown in Fries3 for 72 hrs were left without shaking for a further 14-21 days. The liquid content was then filtered through a sterile 0.22 µm sterile filter before use in leaf infiltrations. Seedlings were returned to the growth chamber (Conviron) and lesions were visually assessed after four days for a qualitative comparison of the necrosis-inducing potential of secretomes for the respective fungal mutants. Five wheat lines were used that are differentially sensitive to ToxA, Tox1 and Tox3 [Halberd (*Tsn1*, *Snn1*, *Snn3*), Calingiri (*tsn1*, *Snn1*, *snn3*), Estoc (*Tsn1*, *snn1*, *Snn3*), BG220 (*tsn1*, *snn1*, *Snn3*) and BG261 (*Tsn1*, *snn1*, *snn3*)].

**Identification of co-expressed transcription factors**

The set of *P*. *nodorum* TFs were sourced from a previous study (21)to specifically identify PnPf2 targets. Their experimentally-validated putative orthologues were compiled based on a literature review (22). A hierarchical-cluster analysis using a genome-wide microarray expression dataset (23) was also undertaken to identify *P*. *nodorum* TFs co-expressed with *PnPf2*, *ToxA*, *Tox1* and *Tox3*. The SN15 genes were clustered together with pheatmap using the normalised microarray gene expression values (Z-scores) (24). Clustering distances were derived from Pearson’s correlation coefficients using the Average linkage function.

**Supplemental text – References**


1. Punt PJ, Oliver RP, Dingemanse MA, Pouwels PH, van den Hondel CA. 1987. Transformation of *Aspergillus* based on the hygromycin B resistance marker from Escherichia coli. Gene 56:117–124.
2. Mattern I, Punt P, Hondel C. 1988. A vector for Aspergillus transformation conferring phleomycin resistance. Fungal Genetics Reports 35:25.
3. Nødvig CS, Nielsen JB, Kogle ME, Mortensen UH. 2015. A CRISPR-Cas9 system for genetic engineering of filamentous fungi. PLoS One 10:e0133085.
4. Sexton AC, Howlett BJ. 2001. Green fluorescent protein as a reporter in the *Brassica–Leptosphaeria maculans* interaction. Physiological and Molecular Plant Pathology 58:13–21.
5. Shaner NC, Campbell RE, Steinbach PA, Giepmans BNG, Palmer AE, Tsien RY. 2004. Improved monomeric red, orange and yellow fluorescent proteins derived from *Discosoma* sp. red fluorescent protein. 12. Nat Biotechnol 22:1567–1572.
6. Engler C, Kandzia R, Marillonnet S. 2008. A one pot, one step, precision cloning method with high throughput capability. PLoS One 3.
7. Andreou AI, Nakayama N. 2018. Mobius assembly: a versatile Golden-Gate framework towards universal DNA assembly. PLOS ONE 13:e0189892.
8. Solomon PS, Tan K-C, Sanchez P, Cooper RM, Oliver RP. 2004. The disruption of a Gα subunit sheds new light on the pathogenicity of *Stagonospora nodorum* on wheat. Mol Plant Microbe Interact 17:456–466.
9. Solomon P, Ipcho S, Hane J, Tan K-C, Oliver R. 2008. A quantitative PCR approach to determine gene copy number. Fungal Genetics Reports 55:5–8.
10. Rybak K, See PT, Phan HTT, Syme RA, Moffat CS, Oliver RP, Tan K-C. 2017. A functionally conserved Zn2Cys6 binuclear cluster transcription factor class regulates necrotrophic effector gene expression and host-specific virulence of two major Pleosporales fungal pathogens of wheat. Mol Plant Pathol 18:420–434.
11. Jones DAB, John E, Rybak K, Phan HTT, Singh KB, Lin S-Y, Solomon PS, Oliver RP, Tan K-C. 2019. A specific fungal transcription factor controls effector gene expression and orchestrates the establishment of the necrotrophic pathogen lifestyle on wheat. 1. Sci Rep 9:1–13.
12. Syme RA, Tan K-C, Hane JK, Dodhia K, Stoll T, Hastie M, Furuki E, Ellwood SR, Williams AH, Tan Y-F, Testa AC, Gorman JJ, Oliver RP. 2016. Comprehensive annotation of the *Parastagonospora nodorum* reference genome using next-generation genomics, transcriptomics and proteogenomics. PLOS ONE 11:e0147221.
13. Teste M-A, Duquenne M, François JM, Parrou J-L. 2009. Validation of reference genes for quantitative expression analysis by real-time RT-PCR in *Saccharomyces cerevisiae*. BMC Mol Biol 10:99.
14. Mead O, Thynne E, Winterberg B, Solomon PS. 2013. Characterising the role of GABA and its metabolism in the wheat pathogen *Stagonospora nodorum*. PLOS ONE 8:e78368.
15. Liu Z, Faris JD, Meinhardt SW, Ali S, Rasmussen JB, Friesen TL. 2004. Genetic and physical mapping of a gene conditioning sensitivity in wheat to a partially purified host-selective toxin produced by *Stagonospora nodorum*. Phytopathology 94:1056–1060.

16. Livak KJ, Schmittgen TD. 2001. Analysis of relative gene expression data using real-time quantitative PCR and the 2(-Delta Delta C(T)) Method. Methods 25:402–408.
17. Solomon PS, Waters ODC, Jörgens CI, Lowe RGT, Rechberger J, Trengove RD, Oliver RP. 2006. Mannitol is required for asexual sporulation in the wheat pathogen *Stagonospora nodorum* (glume blotch). Biochem J 399:231–239.
18. Peters-Haugrud AR, Zhang Z, Richards JK, Friesen TL, Faris JD. 2019. Genetics of variable disease expression conferred by inverse gene-for-gene interactions in the wheat-*Parastagonospora nodorum* pathosystem. Plant Physiology 180:420–434.
19. Solomon PS, Waters ODC, Simmonds J, Cooper RM, Oliver RP. 2005. The Mak2 MAP kinase signal transduction pathway is required for pathogenicity in *Stagonospora nodorum*. Curr Genet 48:60–68.
20. Tan K-C, Phan HTT, Rybak K, John E, Chooi YH, Solomon PS, Oliver RP. 2015. Functional redundancy of necrotrophic effectors – consequences for exploitation for breeding. Front Plant Sci 6.
21. John E, Singh KB, Oliver RP, Tan K-C. 2021. Transcription factor lineages in plant-pathogenic fungi, connecting diversity with fungal virulence. Fungal Genetics and Biology In press.
22. John E, Singh KB, Oliver RP, Tan K-C. 2021. Transcription factor control of virulence in phytopathogenic fungi. Mol Plant Pathol 22:858–881.
23. Ipcho SVS, Hane JK, Antoni EA, Ahren D, Henrissat B, Friesen TL, Solomon PS, Oliver RP. 2012. Transcriptome analysis of *Stagonospora nodorum*: gene models, effectors, metabolism and pantothenate dispensability. Mol Plant Pathol 13:531–545.
24. Kolde R. 2015. pheatmap: Pretty Heatmaps. R package version 1.0.10.
